# Organ-specific microbiomes in natural *Lotus corniculatus* populations: Metacommunity dynamics in the plant endosphere

**DOI:** 10.1101/2025.04.15.649008

**Authors:** Katrina Lutap, Juliana Almario, Maryam Mahmoudi, Frank Reis, Oliver Bossdorf, Eric Kemen

## Abstract

The structure of plant microbial communities vary due to a broad range of factors such as host and environmental factors, abiotic and biotic perturbations, and various assembly processes occurring at multiple tempo-spatial scales. In natural environments plant microbial communities are constantly exposed to such perturbations and processes. Thus, to attain a systemic understanding of the ecology of plant microbiomes, it is essential to study assembly processes that influence patterns of microbial community structures in natural environments. In this study we examined bacterial, fungal, and eukaryotic communities in plant organs of *Lotus corniculatus* in natural populations at seven grassland sites for four years. We used the framework of metacommunity theory of ecology to understand assembly processes that shape community structures and variations by defining microbial communities associated with the roots, shoots, flowers, and seeds as distinct communities linked by dispersal. In this study we show the organ-specificity of plant endophytic communities. Our findings suggest that selective filtering by plant organs, microbial interactions, as well as abiotic and biotic factors at tempo-spatial scales result in distinct core microbiomes of plant organs. In addition, transmission of microorganisms from within and outside the plant hosts accounts for the distinct yet overlapping organ microbiomes. We could provide a comprehensive knowledge of the stochastic and deterministic assembly processes that shape plant microbial communities in natural conditions. Understanding these ecological processes is essential for harnessing beneficial effects of plant-associated microbial communities on plant productivity, resilience, and pathogen defense.

## INTRODUCTION

Plants are associated with diverse microbial communities of beneficial, neutral, and pathogenic microorganisms from across different kingdoms of archaea, bacteria, fungi, protists, and viruses. These plant-microbe interactions can be beneficial for both plant hosts and associated microorganisms - while microorganisms function in plant growth and development, nutrient acquisition, stress tolerance, and pathogen defense, host plants provide habitats and resources for growth and survival of microorganisms (1). These microbes that live within (*i.e*. endophytes) or on the surface (*i.e*. epiphytes) of plants are acquired from seeds via vertical transmission and from biotic and abiotic environments comprising air or wind, rain, soil, insects, and animals via horizontal transmission (2-7). The structure of plant microbial communities vary across plant age and genotypes, geographic locations, soil properties and land use types, developmental stages, and plant compartments (8-12). While many studies on plant microbiomes described these variations in microbial community structures, there remains a need for comprehensive understanding on assembly processes and microbial interactions that account for these community patterns. Current research on plant microbiomes revealed complex associations between plant hosts, microorganisms, and environment, however there are remaining challenges on our understanding of mechanisms that govern community assembly, community structure variations, and intricate interactions in plant microbial communities, especially in natural conditions. Given the high diversity and complexity of microbial communities in their natural habitats, the ecology of microbiomes *in situ* can only be partially conveyed by *in vitro* studies under laboratory conditions (13). To address these challenges, it is crucial to study plant microbiomes in natural environments, with constant exposure to natural perturbations and interactions, biotic and abiotic elements, and natural microbial sources.

Plant microbiomes are shaped by factors acting at multiple scales - from populations to individual plants and plant organs, across time and space (14, 15). Plant microbial communities are also interconnected via transmission of microbes at different levels, from microbial sources outside plant hosts (*i.e* other plants, animals, and environment) to transmission within plant tissues (16, 17). To incorporate simultaneously the community assembly processes occurring at multiple spatial and temporal scales and consequently to attain a more complete understanding of the variations and interactions in plant microbial communities, we examined community dynamics of plant microbiomes in the framework of metacommunity theory of ecology. Metacommunity theory is the study of distinct communities that are linked by dispersal (18). This concept consolidates the effect of different ecological processes and their interactions across space and time scales on community structures (19, 20). The assembly processes defined in Vellend’s conceptual synthesis of community ecology are integrated into the metacommunity theory - selection, drift, and diversification are shaping separately each of the distinct communities, which are linked by dispersal (21, 22). Studies on microbial communities in the context of metacommunity ecology are relatively recent. Traditionally, metacommunity theory is used in the study of spatially distinct macrobial communities, such as marine and freshwater habitats, forest communities, moss patches, and insect and aquatic invertebrate metacommunities (23, 24). Application of the metacommunity concept in microbiome research allows for synthesis of both the deterministic and stochastic processes in their roles on shaping microbial communities at multiple temporal and spatial scales. Microbial communities in different ecosystems or sites (*i.e*. soil and freshwater ecosystems), as well as in plant and animal populations, were analyzed as metacommunities (25-29). Research on host-microbiome systems as metacommunities accounts for the interactions of assembly processes and microbial routes within hosts and their outside environments. A few studies on host-microbiomes and their interactions with their environments have been conducted with communities of insects, aquatic and terrestrial animals, and humans (30-36). Studying plant microbiomes as metacommunities takes into account varied plant host characteristics as well as diverse environmental factors and microbial sources influencing at scales ranging from individual plants to populations. Plant-microbiome systems as metacommunities is a fairly new concept (37). Moreover, studies focusing on microbial communities in plant organs in the context of metacommunity ecology are currently few. Investigation of microbial communities associated with roots, flowers, and seeds as discrete communities interacting with their biotic and abiotic environment were recently presented (38-41).

Studying plant organ-associated microbial communities as distinct communities linked with other microbial communities within and outside the plant host holds advantages towards better understanding of complex assembly processes and microbial interactions. Factors influencing the interconnected microbial communities at multiple tempo-spatial scales - from plant populations to individual plants and tissues - are simultaneously integrated in one ecological framework. In the metacommunity context, the plant host encompasses heterogeneous habitats for microbes, in which each of the plant organs are influenced by different assembly processes, biotic interactions, and microbial transmission, shaping distinct plant organ-specific microbial communities (1, 42, 43). Different resources or microhabitat conditions can cause varied growth and survival responses of microorganisms within plant organs, resulting in organ-specific selection for or against a set of microorganisms (14, 17). The abundance-persistence concepts in ecology can be utilized to determine well-adapted microorganisms consistently interacting with host organs across time and space (22, 44-46). Aside from these host-microbe interactions, distinct microbe-microbe interactions can emerge from compartment-specific environments (47, 48). These microbe-microbe interactions can be inferred through correlation networks and consequently microbial hubs that potentially shape microbial community structures can be identified (49-51). Biotic and abiotic factors acting at tempo-spatial scales ranging from plant organs to populations are simultaneously influencing plant microbiomes (14, 16, 17, 52, 53). By inspecting these biotic and abiotic elements at the level of microbial scales, individual plant scales, to population scales (*i.e*. at levels of plant organs, below- and above-ground, and between sites, respectively), we can identify factors that significantly contribute to observed community patterns in plant organ microbiomes. While plant organ microbiomes are distinct, they are interconnected via transmission of microorganisms among plant organs and from outside of plant hosts (1, 14-17, 53). Dispersal events can be inferred by estimating potential microbial sources of plant organ microbiomes (54). The influence of dispersal on microbial communities via priority effects can also be explored by predicting candidate early-arriving microbes which altered community composition (55).

In this study, we investigated the diversity and community composition of microbiomes associated with *Lotus corniculatus* in natural populations. *L. corniculatus*, a common legume species that is widespread in European grasslands, is an ideal system to study assembly processes and variations of microbial communities in natural environments. *L. corniculatus* is a perennial flowering plant that can adapt to a broad range of natural environments - it can grow on different types of soils, is resistant to grazing and mowing, and is host to various insects and bees (56-58). *L. corniculatus* is also known to host diverse microbes, including nitrogen-fixing rhizobia bacteria and arbuscular mycorrhiza fungi which enhance the plant host’s ability to adapt in poor habitats, as well as non-rhizobial endophytes that promote plant growth (59-61). In this study, we also examined the assembly processes that account for the community structures and variations in *L. corniculatus*-associated endophytic communities in the framework of metacommunity theory in ecology by defining the microbial communities associated with roots, shoots, flowers, and seeds as distinct communities linked by transmission of microorganisms from within and outside the plant hosts. Specifically, we aimed (i) to establish that plant organs host distinct microbial communities by using diversity measures and predictive models in machine learning (*i.e*. organ-specificity of plant microbial communities); (ii) to identify well-adapted microorganisms selectively filtered by each host plant organ by utilizing abundance-persistence concepts in ecology and machine learning approaches (*i.e*. host selection); (iii) to infer important microbe-microbe interactions in plant organs by constructing correlation networks and predicting hubs in the organ-specific microbial communities (*i.e*. microbial interactions); (iv) to explore at different levels of tempo-spatial scales the factors that contribute to plant organ-specific patterns of microbial communities by using diversity measures across sampling years and sites (*i.e*. biotic and abiotic factors); and finally (v) to predict dispersal events that link distinct plant organ communities by estimating transmission of microorganisms from various microbial sources and predicting microorganisms that potentially have roles in priority effects phenomena (*i.e*. microbial transmission and priority effects). To address these aims, we collected *L. corniculatus* from seven grassland sites in the Swabian Alps, Germany for four years. We performed amplicon sequencing of microbial 16S rRNA, ITS2, and 18S rRNA genes targeting endophytic communities in plant organs. Results are synthesized in the framework of the metacommunity concept to establish a comprehensive understanding of ecological processes that shape plant microbial community dynamics.

## METHODS

### Collection of *L. corniculatus* samples in Swabian Alps

To study the ecology of plant microbiomes in natural environments, we collected *Lotus corniculatus* from seven grassland sites in the region of Swabian Alps, Germany for four years (Fig. 4a). These grassland sites of the Biodiversity Exploratories project in southwest Germany have different land use types including unfertilized, mown pastures (AEG3, AEG8, AEG43), fertilized, mown pastures (AEG10, AEG40), and fertilized, mown meadows (AEG17, AEG22) (62-65). We collected plant samples every August-September from year 2018 to 2021 when plants are flowering and producing fruits. We sampled from each site six plants that were randomly distributed throughout the area. We also sampled soil from where plants were uprooted and then pooled together for each site. To characterize endophytic communities in *L. corniculatus* organs, we separated plant samples into roots, shoots, flowers, and seeds and then surface-sterilized them sequentially with sterile water, epiphyte wash (1X TE + 0.1% Triton X-100), 80% ethanol, bleach (2% NaOCl), and finally sterile water. We stored sterilized samples at -20 ^°^C until processing for DNA extraction.

### Amplicon sequencing of *L. corniculatus*-associated bacteria, fungi, and eukaryotes

We sequenced a total of 700 samples of soil and *L. corniculatus* roots, shoots, flowers, and seeds. We homogenized frozen samples of soil and surface-sterilized roots, shoots, flowers, and seeds in Precellys 24 Tissue Homogenizer (Bertin Technologies) before we extracted DNA using FastDNA™ Spin Kit for Soil (MP Bio) as described in the manufacturer’s protocol. Extracted DNA, along with blank samples (*i.e*. water and blank DNA extraction), were used as templates for two-step PCR amplification of bacterial 16S rRNA V5-V7 region, fungal ITS2 region, and eukaryotic 18S rRNA V9 region using primers 799F/1192R, fITS7/ITS4, and F1422/R1797, respectively (Table S1) (51). We designed blocking oligos using R package “AmpStop” to minimize amplification of mitochondrial and chloroplast 16S rRNA, ITS, and 18S rRNA from *L. corniculatus* (Table S1) (66). Amplification products randomized in eight sequencing batches were pooled in equimolar concentrations and purified via magnetic bead clean-up before sequencing on Illumina MiSeq with PhiX control using MiSeq Reagent Kit v3 (600-cycle).

### Sequence data processing

We processed amplicon sequence data of microbial 16S rRNA, ITS2, and 18S rRNA using Mothur as described in Almario *et al*. (Method S1) (52, 67). We taxonomically classified bacterial 16S rRNA, fungal ITS2, and eukaryotic 18S rRNA sequences based on Greengenes database (13_8_99 release), UNITE database (02.02.2019 release), and PR^2^ database (version 4.12.0), respectively, with the PhiX genome included in the databases (68-70). For 16S rRNA and 18S rRNA data, Cutadapt was used to remove primer sequences, and for ITS2 data, ITSx was used to remove non-ITS sequences (71, 72).

### Diversity and community composition analysis

For diversity analysis and relative abundance calculations, we used R packages phyloseq, vegan, microbiome, and microeco to analyze OTU tables outputted from Mothur pipeline (73-76). We used Shannon’s diversity and Observed species indices to assess alpha-diversity of samples. To check if alpha-diversity measures between samples are significantly different, we tested data whether they are normally distributed via Shapiro-Wilk normality tests and then analyzed using parametric test ANOVA (for normally distributed data) or nonparametric Kruskal-Wallis rank sum test (for non-normal data). We conducted post-hoc analysis using Dunn’s test (for Kruskal-Wallis test) or Tukey’s HSD (for ANOVA test). For beta-diversity analyses, we used OTU relative abundance tables for Principal Coordinate Analysis (PCoA) ordination of Bray-Curtis dissimilarities between samples. We used PERMANOVA analysis of Bray-Curtis distances to assess significant explanatory variables (*i.e*. plant organ, years of sampling, sampling sites) affecting microbial community structures.

### Identification of key organ-specific microbes in *L. corniculatus*

To determine abundant endophytes in *L. corniculatus* roots, shoots, flowers, and seeds, we identified OTUs with highest relative abundances in each organ (≥1% relative abundance). To determine persistent core microbes in organs, we identified OTUs that occur in at least 90% of the samples across seven sampling sites for four years (≥90% occurrence). To determine hub bacteria, fungi, and eukaryotes in plant organs, we computed correlation networks for each plant organ using SparCC algorithm, as described in Almario *et al*. (Method S2) (52, 77). We used Cytoscape (v.3.9.1) to visualize networks and to calculate network features such as number of nodes and edges, node connectedness (degree), betweenness centrality, and closeness centrality (78). We assigned OTUs that are top 5% in betweenness centrality and closeness centrality scores as hub microbes. We used representative sequences of abundant, core, or hub OTUs for multiple sequence alignment in MUSCLE (Web Form) using default parameters (79). We used resulting alignments in ClustalW format to build neighbor-joining phylogenetic trees in the online tool iTOL (v.6.7.3) (80).

### Machine learning classification for diagnosing multi-organ involvement and predicting organ-specific microbes in *L. corniculatus*

To explore the possibility of a predictive pattern for separation of *L. corniculatus* organs based on microbial composition, we employed a linear classification machine learning model using relative abundance data of bacteria, fungi, and eukaryotes. We utilized one-vs-one approach on the support vector machine with a linear kernel (OneVsOneClassifier(svm.SVC) function in Scikit-learn) to train a binary classification model for separating each organ from others (81). Using Scikit-learn, we randomly selected 67% of samples (n=467) for training and reserved the remaining 33% (n=231) for independent testing and evaluation using the classification_report function, then we visualized confusion matrices using the plot_confusion_matrix function. To identify microbial species that can discriminate each organ from the rest, we employed recursive feature elimination with cross-validation (RFECV function). We calculated the accuracy of the model using K-fold cross-validation (K=10).

### Transmission of microbes in *L. corniculatus*

To determine how *L. corniculatus* organ microbiomes can be linked and influenced by dispersal, we used Sankey diagrams to visualize potential flow of microbes across the soil and plant organs (Method S3). To determine potential origins of organ-associated microbial communities, we used FEAST (Fast Expectation-mAximization microbial Source Tracking) to estimate contribution of potential microbial sources, such as soil, the different plant compartments, or the environment, to each plant organ microbiome (Method S3) (54). To statistically predict potential priority effects phenomena from roots, shoots, flowers, to seeds in *L. corniculatus*, we identified taxa of interest that are potentially involved in such phenomena in plant organs, as described in Debray *et al*. (Method S3) (55).

## RESULTS AND DISCUSSION

### Endophytic bacterial, fungal, and eukaryotic communities in natural *L. corniculatus* populations are organ-specific

We surveyed endophytic communities associated with *L. corniculatus* in natural populations by amplicon sequencing of bacteria, fungi, and eukaryotes in 700 samples of soil and plant roots, shoots, flowers, and seeds (Table S2, Table S3). We used blocking oligos which decreased the number of nontarget plant DNA reads (*i.e*. chloroplast, mitochondria, plant ITS2, and plant 18S rRNA) by 90-100% and significantly increased the number of microbial 16S rRNA, ITS2, and 18S rRNA reads (Fig. S1). After processing raw reads and removing unknown and plant sequences, we identified a total of 4,225 16S rRNA, 2,027 ITS2, and 1,773 18S rRNA OTUs clustered based on 97% sequence similarity and were classified into 113 phyla and 1,542 genera (Table S3). Among the plant compartments, roots have the greatest number of OTUs followed by shoots, while there are less number of OTUs detected in flowers and in seeds (Fig. S2a). In all plant compartments, phyla Proteobacteria, Actinobacteria, Firmicutes, Bacteroidetes, Acidobacteria, Ascomycota, Basidiomycota, and unclassified Fungi and Eukaryota are the most abundant groups (Fig. 1a). In addition, in roots phyla Chloroflexi and Nematoda are abundant, while in shoots, flowers, and seeds, Arthropoda and unclassified Bacteria dominate. Soil samples have more observed OTUs compared with all plant organs, with groups Actinobacteria, Proteobacteria, Ascomycota, Basidiomycota, and unclassified Fungi as the most abundant phyla (Fig. 1a, Fig. S2a).

**Figure 1.**
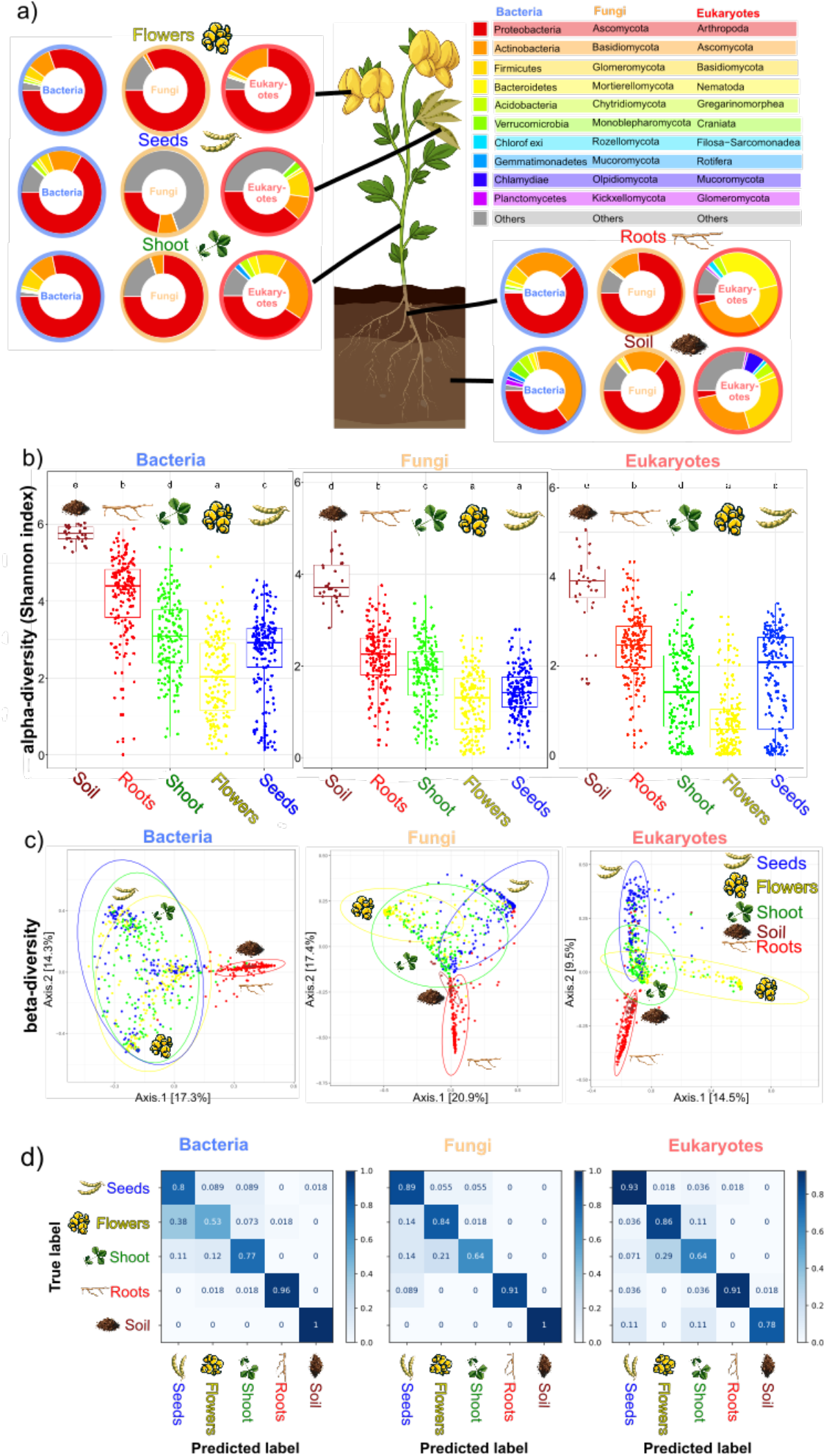
Organ-specificity of plant microbial communities. **(a)** Relative abundance profiles showing the top ten most abundant bacterial, fungal, and eukaryotic phyla in soil and plant organ samples collected from seven grassland sites for four years. The relative abundance of the OTUs for each compartment (*i.e*. soil and plant organs) were aggregated at the phylum level. **(b)** Boxplots of Shannon’s α-diversity measurements of bacterial, fungal, and eukaryotic microbial communities associated with soil and plant organs. Shapiro-Wilk normality tests indicated that datasets have non-normal distribution (p<0.05) and Kruskal-Wallis rank sum tests were used to test significant differences of α-diversity measurements between soil and plant organ samples. Post-hoc analysis via Dunn’s tests indicate that groups are significantly different if letters are not similar (Supplementary Table 4). **(c)** Principal coordinate plots based on Bray-Curtis dissimilarities between bacterial, fungal, and eukaryotic microbial communities associated with all soil and plant organ samples. **(d)** Performance of the multi-class support vector machine (SVM) model on the test set showing high accuracy in separating bacterial, fungal, and eukaryotic communities in organs and soil samples.

Alpha-diversity analysis showed that endophytic communities in roots are the most diverse compared with communities in aboveground plant organs. Shannon diversity index showed that endophytic communities progressively become less diverse from roots to shoots and flowers, and then an increase in bacterial and eukaryotic diversity is observed in seeds (Fig. 1b, Table S4b). Soil microbial communities are more diverse than microbial communities associated with *L. corniculatus* (Fig. 1b, Table S4b). Beta-diversity analysis revealed that the diverse and stable microbial communities in soil and roots are distinct from the less diverse and more variable communities in shoots, flowers, and seeds. Bray-Curtis dissimilarities-based PCoA showed that soil and root microbial communities are similar in community structures and are distinct from the overlapping endophytic communities in shoots, flowers, and seeds (Fig.1c). The PCoA plots showed a main separation (axis 1) of bacterial communities between above and belowground communities while fungal and eukaryotic communities separate the aboveground organs. Both PCoA and relative abundance profiles of individual samples across sampling sites and years also showed that soil and root microbial communities are more clustered and less dispersed compared with the variable microbial communities in shoots, flowers, and seeds (Fig. S3). PERMANOVA indicated that significant separation of microbial communities was largely explained by plant organ, while sampling years and sites contributed relatively lower to microbial community variations (Fig. S4; Table S4c).

We trained a multi-class support vector machine (SVM) model to determine whether relative abundances of bacterial, fungal, and eukaryotic microbial communities could distinguish between each organ from other organs (all binary possibilities) and soil samples. The performance of the model on the test set showed high accuracy in separating organs and soil samples (accuracy = 77-83%; Table S5). In bacterial and fungal communities, roots and soil samples separated from other groups with higher prediction accuracy compared with aboveground organs, while in eukaryotic communities, seeds are most accurately predicted among all the groups (Fig. 1d). We trained an SVM classifier with recursive feature elimination and cross-validation to identify OTUs that discriminate each organ from the others (without soil samples). Results revealed a subset of 57-166 OTUs that could separate roots, shoots, flowers, and seeds from other organs (Fig. S5).

Based on diversity analyses and predictive models, results show the organ-specificity of endophytic communities in natural *L. corniculatus* populations. Root microbial communities are the most diverse among the plant organ communities. There is significant separation of community structures between aboveground and belowground bacterial communities, while shoot, flower, and seed communities are mainly separated by fungi and eukaryotes. The aboveground endophytic communities are more variable compared with the root communities. Soil microbial communities, which are more diverse than all plant organ communities, have overlapping community composition with the root communities.

### *L. corniculatus* core microbiomes are shaped by host selection

To identify key microbes which can explain the identified organ-specific patterns in community structures of *L. corniculatus* microbiomes, we examined species abundance, an ecological pattern that is observed in macrobial and microbial communities (Fig. 2, Table S6). Except for *Pantoea* and *Pseudomonas*, which are abundant in both above- and below-ground plant organs, the most abundant OTUs (relative abundance (RA) > 1%) in root microbial communities are distinct from the set of abundant OTUs in aboveground microbial communities. Similarly in microbial communities of shoots, flowers, and seeds, the most abundant OTUs (RA > 1% in at least one plant organ) are mostly the same and enriched specifically in the aboveground microbial communities. The observed similarity in the sets of abundant taxa accounted for the observed overlapping community structures of the aboveground plant organ communities despite high variability throughout sampling years and sites (Fig.1c, Fig. S3). Thus, the overlapping community structure in shoots, flowers, and seeds, as well as the distinction between aboveground and belowground plant organ community structure, can be largely explained by the most abundant species.

**Figure 2.**
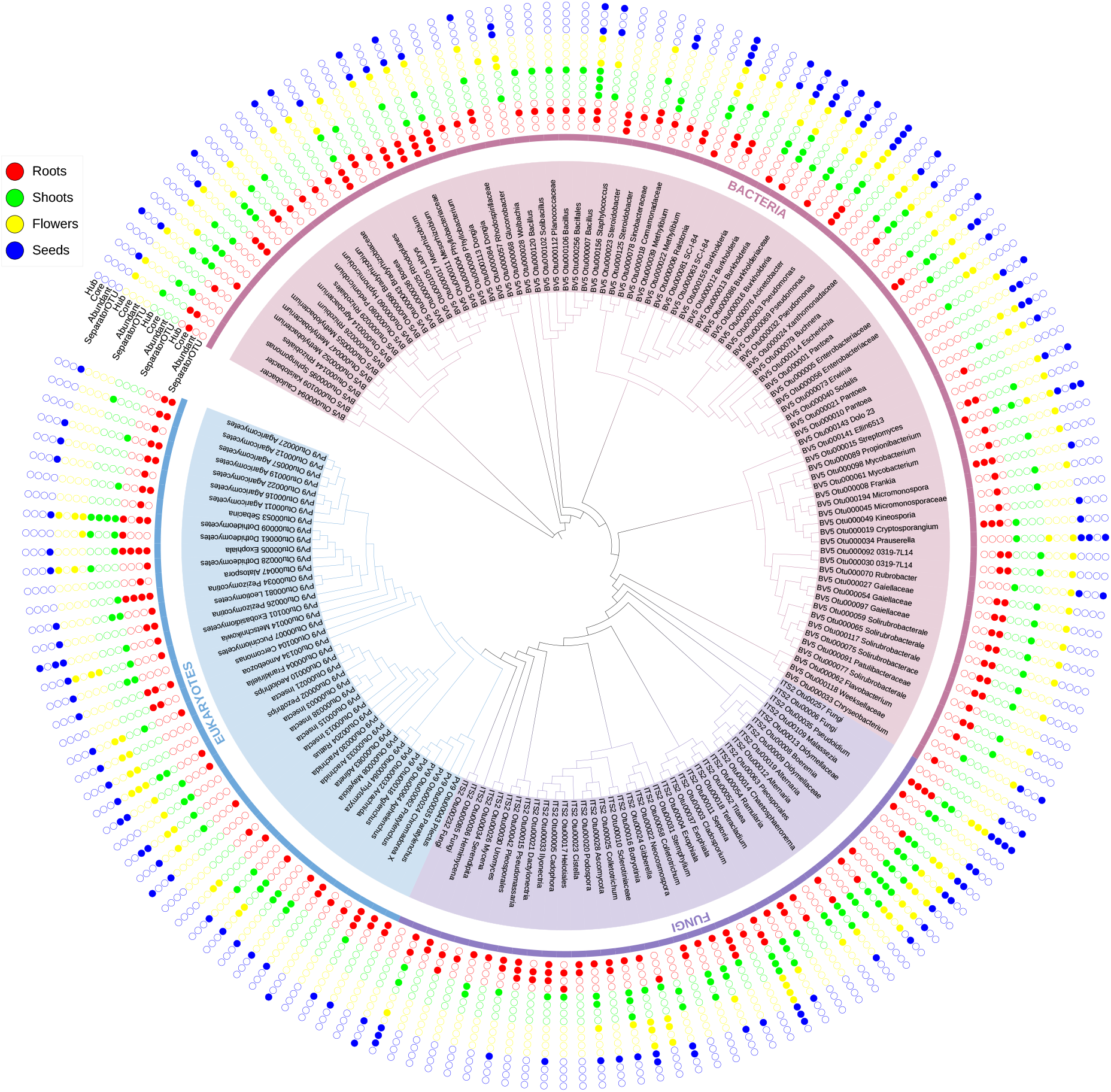
Organ-specific microbes identified via abundance, persistence, and predictive machine learning models. Neighbor-joining phylogenetic tree of top abundant OTUs (relative abundance in the plant organ microbial communities > 1%) and core OTUs (detected in 90% of the organ samples). Abundant and core OTUs that are either separator OTUs (all identified separator OTUs using multi-class support vector machine (SVM) model are detailed in Supplementary Fig. 5) or hubs (hubs identified using microbial networks are detailed in Fig. 3 and Supplementary Fig. 6) are also indicated.

Persistence of species over time and space is also an ecological pattern that is important in structuring macrobial and microbial communities. We identified persistent organ-specific ‘core’ microbes that can be detected in 90% of *L. corniculatus* organ samples collected across all seven sites and four years (Fig. 2, Table S7). In roots we found 56 bacterial, seven fungal, and seven eukaryotic core OTUs, which consists 1% of all 7,504 root OTUs. Many of the core species identified in roots are also abundant species in roots. In shoots there are seven bacterial, four fungal, and four eukaryotic core OTUs (0.3% of 5,223 total shoot OTUs), while in flowers and seeds there are seven (0.2% of 3,706 total flower OTUs) and eight (0.3% of 3,706 total seed OTUs) core OTUs, respectively. Likewise, most of the core species in aboveground communities are also abundant in their respective plant organ microbial communities. The smaller core communities in shoots, flowers, and seeds suggest fluctuating aboveground microbial communities, in contrast with the more stable root communities with a relatively larger core community. Meanwhile, *Pseudomonas* and *Cladosporium* are persistent over time and space in all plant organs, suggesting stable associations and high adaptation to the plant endophytic environment.

To determine microbes that distinguish between *L. corniculatus* organs, we used multi-class support vector machine (SVM) model that predicts OTUs that are highly associated with each organ (Fig. S5). In root microbial communities, 84 separator OTUs were identified with the root compartment. Among these OTUs are abundant species in roots and are persistent throughout years and sites of collection (Fig. 2, Table S7). *Pseudomonas, Phyllobacterium, Frankia, Mesorhizobium, Cryptosporangium, Steroidobacter, Rhizobium*, and *Bosea* are abundant and core bacteria that showed high association with roots. Abundant and core fungi such as *Exophiala, Cadophora*, and *Dactylonectria* are also distinctive of root microbial communities. In shoot microbial communities there are 102 separator OTUs identified, among these OTUs are core and abundant species *Bacillus, Agrobacterium* and *Alternaria* (Fig. 2, Table S7). There are 57 OTUs that are highly associated with the flower compartment including abundant and core fungi *Cladosporium* and insects *Pezothrips* (Fig. 2, Table S7). There are 166 OTUs that are distinctive of seed microbial communities (Fig. 2, Table S7). *Pseudomonas, Ralstonia*, and *Cladosporium* are abundant and persistent species that are highly associated with seeds. Ralstonia are known pathogens of various plants (Hayward 1991). Different resources or habitat conditions in plant organs could result in the filtering of different groups of microorganisms adapted to organ-specific features, consequently resulting in organ-specific distinctive microbial communities (82-94). These microorganisms that are selectively filtered by each of the plant organs are likely specialized microbes that are able to thrive in the unique physical microstructures of the organs while improving plant growth, nutrient uptake, and resistance to stress and diseases They could supply nutrients while in turn benefitting from resources in particular organs. Alternatively they could be pathogens that hijack the host’s genetic resources.

### Community structures of *L. corniculatus* microbiomes are influenced by microbial interactions

Species interactions are also important in maintaining structures of macrobial and microbial communities. Microbial species uniquely adapted to a particular organ environment that can significantly affect microbial interactions consequently can also affect organ-specific microbial community structuring. To infer potential microbe-microbe interactions and to identify hubs in the plant organ communities, we built correlation networks based on species abundances (Fig. 3). In general, root microbial community networks are more complex than networks of aboveground organs. In addition, bacterial networks are more complex compared with fungal and eukaryotic networks in all plant organs. The number of nodes are consistently highest in roots in all bacterial, fungal, and eukaryotic networks, followed by microbial community networks of shoots, flowers, and seeds. (Fig. 3b). Root networks also have the highest number of edges, which correspond to significant correlations between microbes, compared with aboveground organ networks (Fig. 3c). Based on ANOVA and Tukey’s HSD, the nodes of root microbiome networks have the highest number of interactions among all the organ microbiome networks (Fig. 3d). Aboveground, shoot microbiome networks have higher node connectedness compared with flower and seed networks, which have node connectedness that are not significantly different. There are more nodes that are shared between shoots and flowers in bacterial and fungal networks compared with other organ-to-organ networks, while in eukaryotic networks the root-to-shoot networks have the highest number of shared nodes (Fig. 3e).

**Figure 3.**
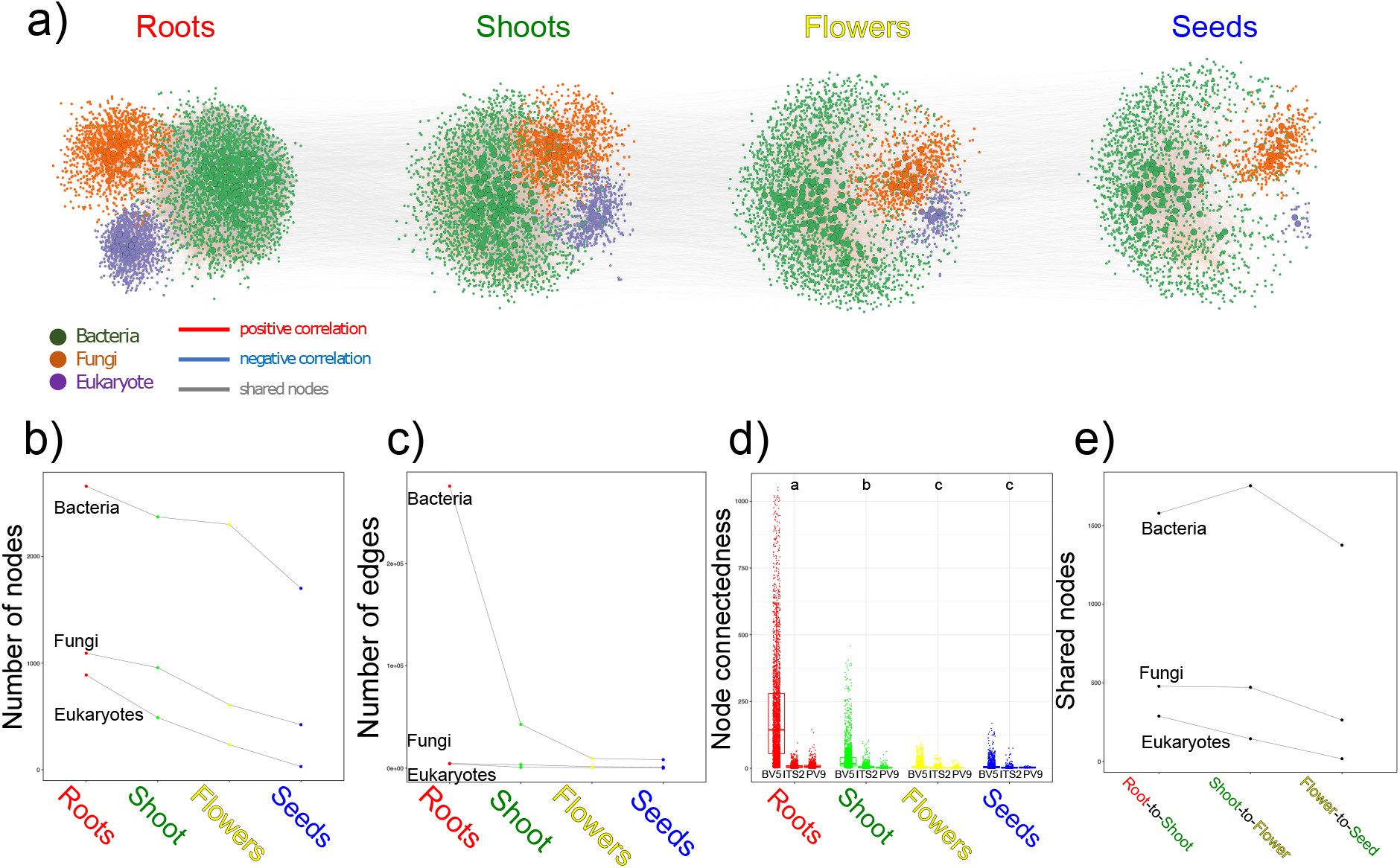
Microbial interactions in organ-specific plant microbiomes. **(a)** Correlation networks based on species abundance of bacterial (green), fungal (orange), and eukaryotic (purple) microbes of each plant organ. Nodes are OTUs and edges represent potential microbe-microbe interactions. Significant positive interactions are colored red, while negative interactions are blue (P<0.001). Hub microbes of each organ microbial communities are represented by bigger and bordered nodes. Grey lines that connect each organ microbiome network show connections of nodes/OTUs that are shared with the next organ. Network characteristics such as number of **(b)** nodes and **(c)** edges, **(d)** node connectedness (degree), and **(e)** nodes shared between organs are used to describe the networks. Overall **(d)** node connectedness (node degree of all bacterial (BV5), fungal (ITS2), and eukaryotic (PV9) networks) were compared between organ microbiome networks via ANOVA and Tukey’s HSD.

Hub microbes, which are highly connected with other microbes in the network, potentially have essential roles in plant-microbe and microbe-microbe interactions and thus shape the community structure (51). We identified hub microbes which can significantly influence microbial community structures in each plant organ based on betweenness centrality and closeness centrality scores (Fig. 2, Fig. S6, Table S7). Many among the hub microbes are abundant and persistent across seven sampling sites throughout four years. In root microbial communities there are 101 bacterial, 39 fungal, and 32 eukaryotic hubs, which comprise 2% of total root OTUs. Many of the inferred hubs are also machine learning-predicted separator microbes in roots, including the hub fungi *Exophiala* that are both abundant and persistent species in roots. Shoot microbial communities comprise 97 bacterial, 33 fungal, and 13 eukaryotic hubs (3% of all shoot OTUs). Separator microbes for the shoot compartment *Bacillus, Alternaria*, and Dothideomycetes are hub microbes that are both persistent and abundant in shoots. Flower-associated microbial communities have 90 bacterial, 23 fungal, and 8 eukaryotic hubs, which consists 3% of total flower OTUs. Expectedly, most of the hubs are separator microbes, including *Cladosporium*, species that are also abundant and persistent in flower compartments. In seed microbial communities, there are 61 bacterial, 14 fungal, and 4 eukaryotic hubs, which is 2% of total seed OTUs. *Pseudomonas* and *Ralstonia* are hub microbes that are both core, abundant, and seed separator microbes. Among the predicted hubs in the plant organs are microbes that are abundant, or persistent across sampling years and sites, and highly associated with the respective organ compartments and are thus important in diversity, stability, and organ-specificity of the microbial communities. These hub microbes either as pathogens or beneficial endophytes play important roles in microbial interactions and can be important in shaping microbial communities of each plant organ (84, 88-94).

### Variation in *L. corniculatus* microbiomes are driven by biotic and abiotic factors across time and space

Beta-diversity analysis of microbiomes in *L. corniculatus* natural populations revealed that plant organs largely contributed to microbial community variation, while years of collection and sampling sites showed smaller effects (Fig.1c, Fig. S4, Table S4c). To further dissect other factors that can influence community structure in *L. corniculatus* microbiomes, we analysed diversity and community composition on each plant organ across sampling years and sites. Throughout sampling sites and years, *L. corniculatus* plants as a whole maintained significantly different levels of microbiome alpha-diversity (except the fungal community), and at least one of the bacterial, fungal, or eukaryotic communities of the plant organs exhibited significantly different diversities (Fig. S7, Fig. 4b-g). Relative abundance profiles of the most abundant bacterial, fungal, and eukaryotic groups in the plants also vary across all sampling sites and years (Fig. S8). Specifically, in roots the diversity of microbial communities is similar across all sampling plots, and while the diversity of fungal and eukaryotic communities are also similar across all sampling years, root bacterial communities exhibited different alpha-diversities throughout the years (Fig. 4b-g). In contrast, microbial communities in aboveground plant organs have significantly different alpha-diversities throughout the years, except in fungal communities of all aboveground organs and in shoot eukaryotic communities (Fig. 4e-g). At least one of the bacterial or fungal communities associated with shoots, flowers, and seeds showed significantly different alpha-diversities across different sampling sites, while eukaryotic communities are similar (Fig. 4b-d).

**Figure 4.**
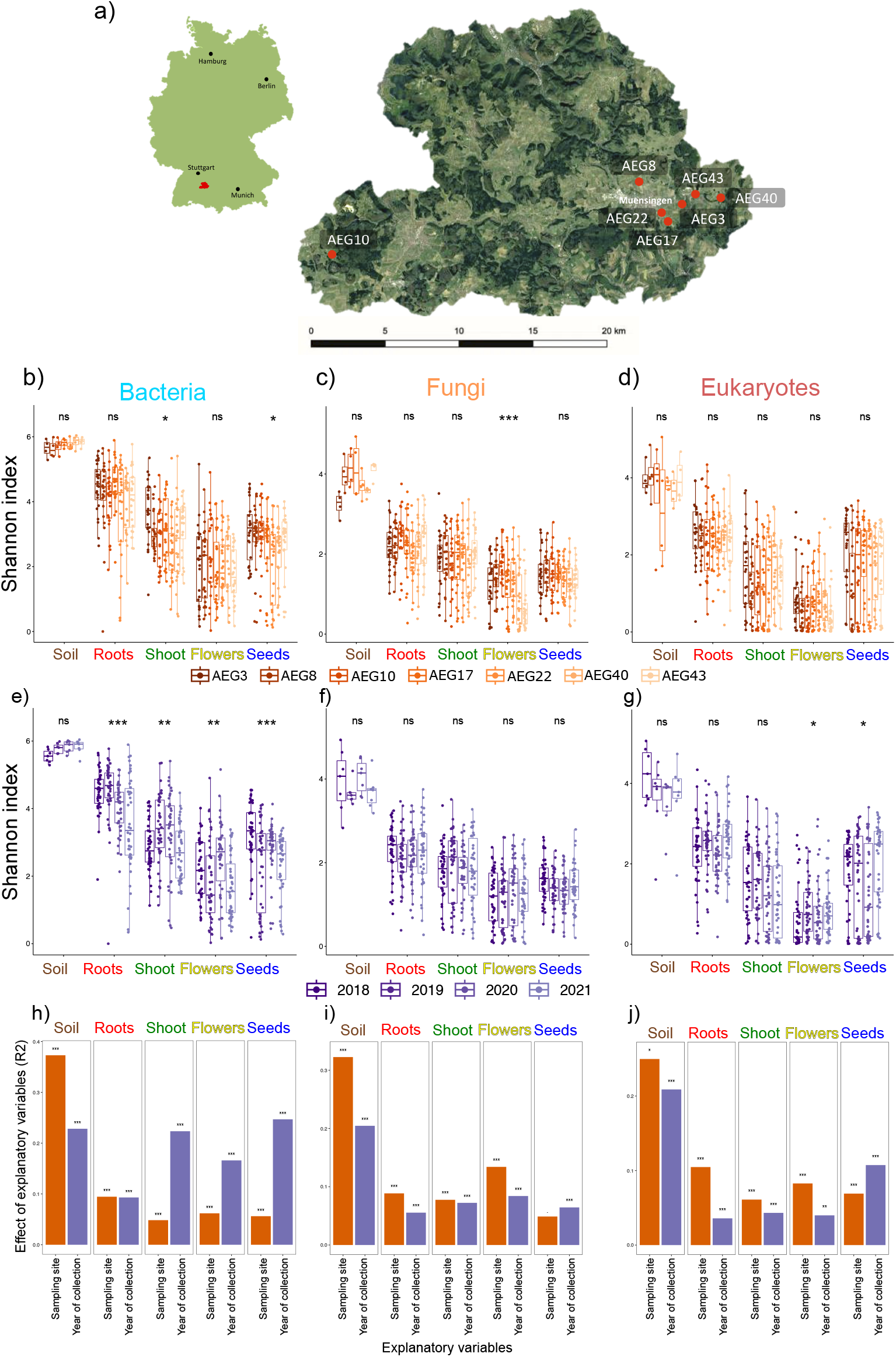
Diversity and community composition of plant organ-associated microbiomes in *Lotus corniculatus* populations in seven grassland sites for four years. **(a)** Map of the seven grassland plots in the Swabian Alps, Germany where samples were collected from years 2018-2021. α-diversity (Shannon index) of the bacterial, fungal, and eukaryotic microbial communities associated with soil and plant organs were compared between **(b-d)** sampling sites or **(e-g)** collection years using Kruskal-Wallis significance test. The effect of explanatory variables (*i.e*. sampling sites, year of collection) on the β-diversity of soil- and plant organ-associated microbial communities were also assessed using **(h-j)** PERMANOVA.

Beta-diversity analysis of *L. corniculatus* microbial communities indicated that years of collection accounted more on the observed variation in bacterial communities compared to sampling sites (Fig. S4, Table S4c). Specifically, while in roots both variables have almost equal effects on variation, the influence of years of collection is higher in the aboveground microbial communities (Fig. 4h, Fig. S9a). These observations show that bacterial communities in roots are relatively more stable across all years and sites, while aboveground communities varied through years of collection, consistent with the relative abundance profiles of bacterial communities in plant organs throughout all years and sites (Fig. S3a-b). *L. corniculatus* are perennial plants, hence the roots could have maintained stable bacterial communities over the years, while in aboveground plant organs the communities are more variable due to consistent perturbations in the sampling sites such as mowing or animal grazing. The recurrent emergence of shoots, flowers, and seeds amidst such perturbations potentially contributed to the variation in bacterial communities in these organs (95, 96). On the other hand, fungal and eukaryotic community structures in all plant organs except seeds are influenced to a larger extent by sampling sites (Fig. 4i-j, Fig. S9b-c). The larger effect of sampling sites suggests that local environmental conditions and soil microbial communities of the different grassland sites have larger influence on the variation of fungal and eukaryotic communities associated with *L. corniculatus* roots, shoots, and flowers. Bacterial, fungal, and eukaryotic community variations in the soil are also influenced largely by sampling sites (Fig. 4h-j, Fig. S9), hence soil microbiomes could have contributed to the beta-diversity patterns observed in the plant organs. Disturbances in the local environments like temperature fluctuations, wind, UV levels, and precipitation, and consequently plant adaptations to these changes, can also potentially augment to the variations of fungal and eukaryotic communities in roots, shoots, and flowers. Insect visitors present in the sampling sites, especially pollinators, can also affect microbial community composition in plants (97). Thus, observed patterns of diversity and composition could be attributed to abiotic factors and biotic interactions acting at different spatial and temporal scales, that is, at the level between roots and aboveground plant organs to the level of sampling locations or years.

### *L. corniculatus* microbiomes are distinct communities linked and influenced by dispersal

To investigate how *L. corniculatus* recruits microbes to assemble into organ-specific communities, we compared OTUs detected across soil and plant organs (Fig. 5a). The distinct but overlapping plant organ microbiomes share a subset of their communities. Most OTUs that are present in plant organs are also detected in soil samples. With increasing distance from the soil, soil-detected OTUs in the plant organs decrease (Fig. S10). Only a small proportion of the core communities in roots are transmitted to upper plant organs. Core microbes *Pseudomonas* and *Cladosporium* are potentially good disperser microbes that are consistently transmitted and maintained throughout the interconnected plant organs (Fig. 2). Mostly, the OTUs in plant organs are transmitted from neighboring compartments and a lesser proportion are from other sources, presumably from the local environment. These observations suggest that there are similarities between plant organ microbial communities due to initial colonizer microbes from soil, while variations in community composition are accumulated through various microbial sources from the environment as well as from other plant compartments. Thus, plants establish their organ-specific microbiomes by recruitment and selecting microbes from soil and their environment as well as by transmission of microbes from other plant compartments.

**Figure 5.**
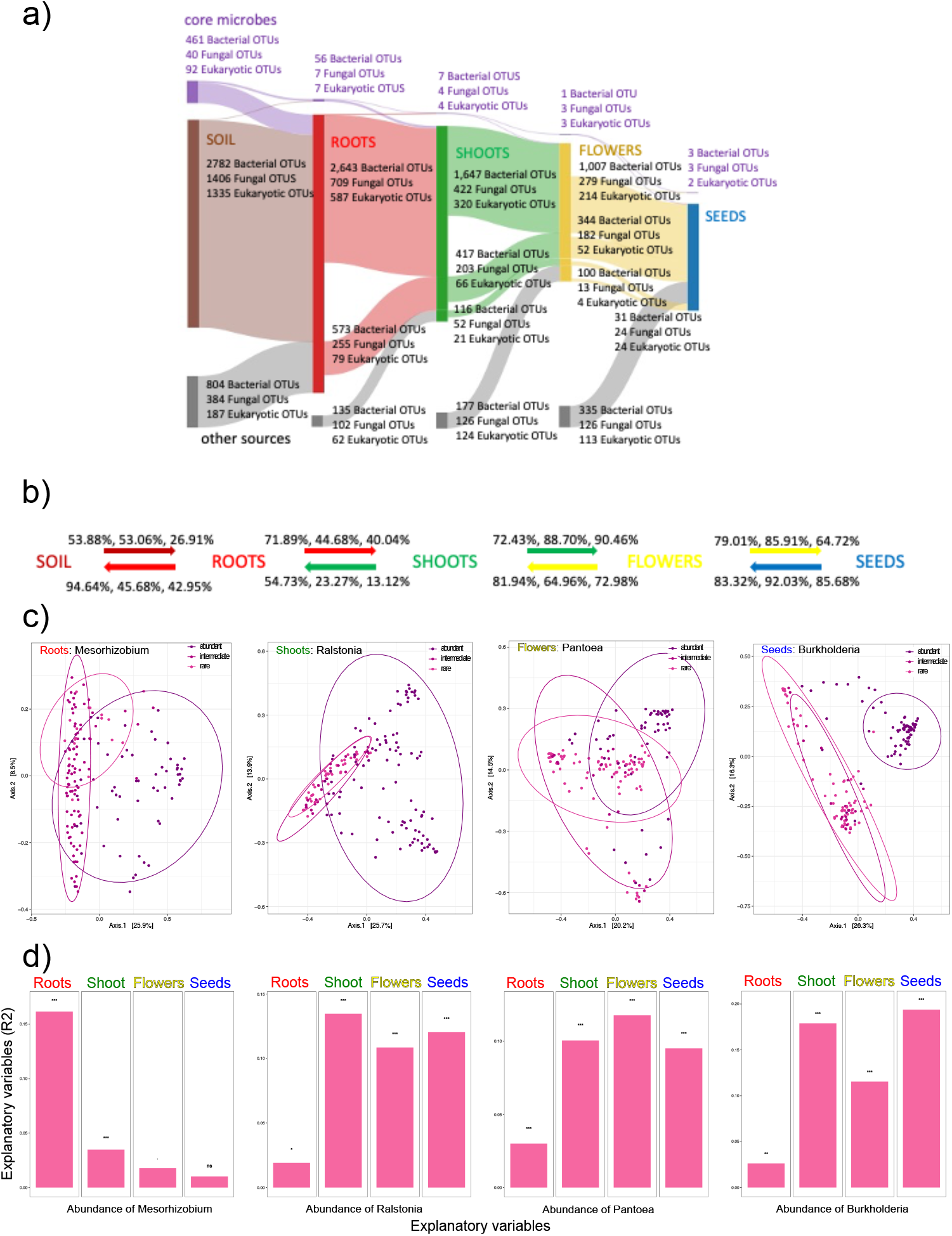
Recruitment of microbes from soil and environment to establish organ-specific microbiomes in *Lotus corniculatus*. **(a)** Sankey diagram of shared microbes between neighboring plant compartments as well as from other sources (*i.e*. from local environment). Nodes of the diagram represent different potential microbial sources (*i.e*. soil, plant organs, and environment/others) and arcs correspond to the number of bacterial, fungal, and eukaryotic OTUs shared between nodes. The number of core OTUs in each plant organ are displayed in purple nodes and arcs in the diagram. **(b)** FEAST was also used to estimate the contribution (% bacterial, fungal, and eukaryotic contribution, respectively) of potential microbial sources to each plant organ microbiome. In FEAST the direction of microbial transmission is preassigned, thus calculation of microbial contribution was tracked at both directions. For more details of transmission at multiple directions, see Supplementary Fig. 11. **(c)** PCoA of Bray-Curtis dissimilarities of plant organ microbial communities and **(d)** PERMANOVA using abundance of candidate early-arriving OTU as explanatory variable of community composition variation. (c) and (d) show that candidate early-arriving OTUs (BV5_OTU11_*Mesorhizobium*, BV5_OTU6_*Ralstonia*, BV5_OTU1_*Pantoea*, BV5_OTU16_*Burkholderia*) potentially altered the microbial community composition of roots, shoots, flowers, and seeds, respectively, based on their abundance in the plant organ microbial communities. (Explanatory variables: abundant OTU: Relative abundance in each plant organ microbial community (RA) >= 0.01; intermediate OTU: RA >= 0.001 and < 0.01; rare OTU: RA < 0.001).

To further verify these observations, we used FEAST to determine potential origins of the organ-specific microbiomes (Fig. 5b). FEAST was used to estimate the contribution of potential microbial sources, such as soil, the different plant compartments, or the environment, to each plant organ microbiome. Since in FEAST the source and sink communities are preassigned and consequently the direction of transmission is not determined, we tracked the transmission at multiple directions (Fig. S11). We observed that there is potential dispersal of microbes from various microbial sources and multiple directions. A large proportion of plant microbes are dispersed between plant compartments (13%-92% of microbes in sink organs are from other plant compartments). Soil microbes are transmitted to all plant organs, and aboveground plant organs tend to have lesser soil microbes compared with roots. Certain fractions of plant organ microbiomes were assigned by FEAST to “unknown sources”, which can be other potential microbial sources from the environment such as insects, pollinators, animals, rain, wind, or soil splashes. These calculations showed that dispersal occurs between plant organ microbiomes as well from soil and environment and thus influences community structures of these organ-specific microbiomes.

Dispersal can also affect microbial community diversity through arrival history. A set of early-arriving species can impact assembly of communities by changing resources or environmental conditions, in a historically-contingent community assembly called priority effects. Debray *et al*. presented an approach to statistically predict from a natural microbiome dataset which taxa are potentially involved in influencing succession during community assembly (55). Based on such approach, we identified candidate early-arriving OTUs that may have inhibited or facilitated establishment of other OTUs by examining if their abundance in plant organs correlated with altered community composition. Changes in relative abundance of some of the key OTUs in plant organs consequently showed changes in community structure (Fig. 5c-d, Fig. S12). For instance, relative abundances of *Phyllobacterium* and *Mesorhizobium* contributed to the variation of microbial communities in roots, while relative abundances of *Pantoea, Ralstonia*, and *Burkholderia* accounted for the microbial community variations in aboveground organs. *Phyllobacterium* and *Mesorhizobium* are abundant, persistent, and highly associated microbes in roots (Fig. 2). *Pantoea* are abundant throughout the whole plant and are core microorganisms in shoots, while *Ralstonia* and *Burkholderia* are key microbes (*i.e*. either abundant, core, hub, or separator microbes) in most of aboveground organs (Fig. 2). The prediction of these taxa and their potential roles during priority effects phenomenon enables future experimental manipulation of arrival history in complex natural microbial communities.

## CONCLUSIONS

We examined the diversity and community composition of *Lotus corniculatus* microbiomes in natural populations in the framework of metacommunity theory to gain broader insight on the ecology of plant microbiomes *in situ*. In this study we showed the organ-specificity of endophytic communities of *L. corniculatus* and established an overview of the assembly processes at tempo-spatial scales that account for the community patterns observed in plant microbiomes in natural populations (Fig. 6). Analysis of community composition and diversity of *L. corniculatus* microbiomes revealed that plant organs are the main source of variation in microbial community structure, while sampling years and sites contributed less. It has been shown that plant compartments contribute more in shaping microbial community composition than geographical locations or sampling times (95, 98, 99). Other studies found that plant compartments account more for associated bacterial community composition, while geographical locations of host plants rather determine fungal community composition (9, 100). Plant organs select for a group of microorganisms that developed adaptive traits to successfully inhabit their unique microenvironments, and in *L. corniculatus* organs abundant and persistent microorganisms are beneficial microbes that enhance growth and fitness, or pathogens that utilize host’s genetic and physiological resources. In the same way, hub microbes in *L. corniculatus* organs are abundant and persistent microbes that are either known plant pathogens or beneficial to plant hosts.

**Figure 6.**
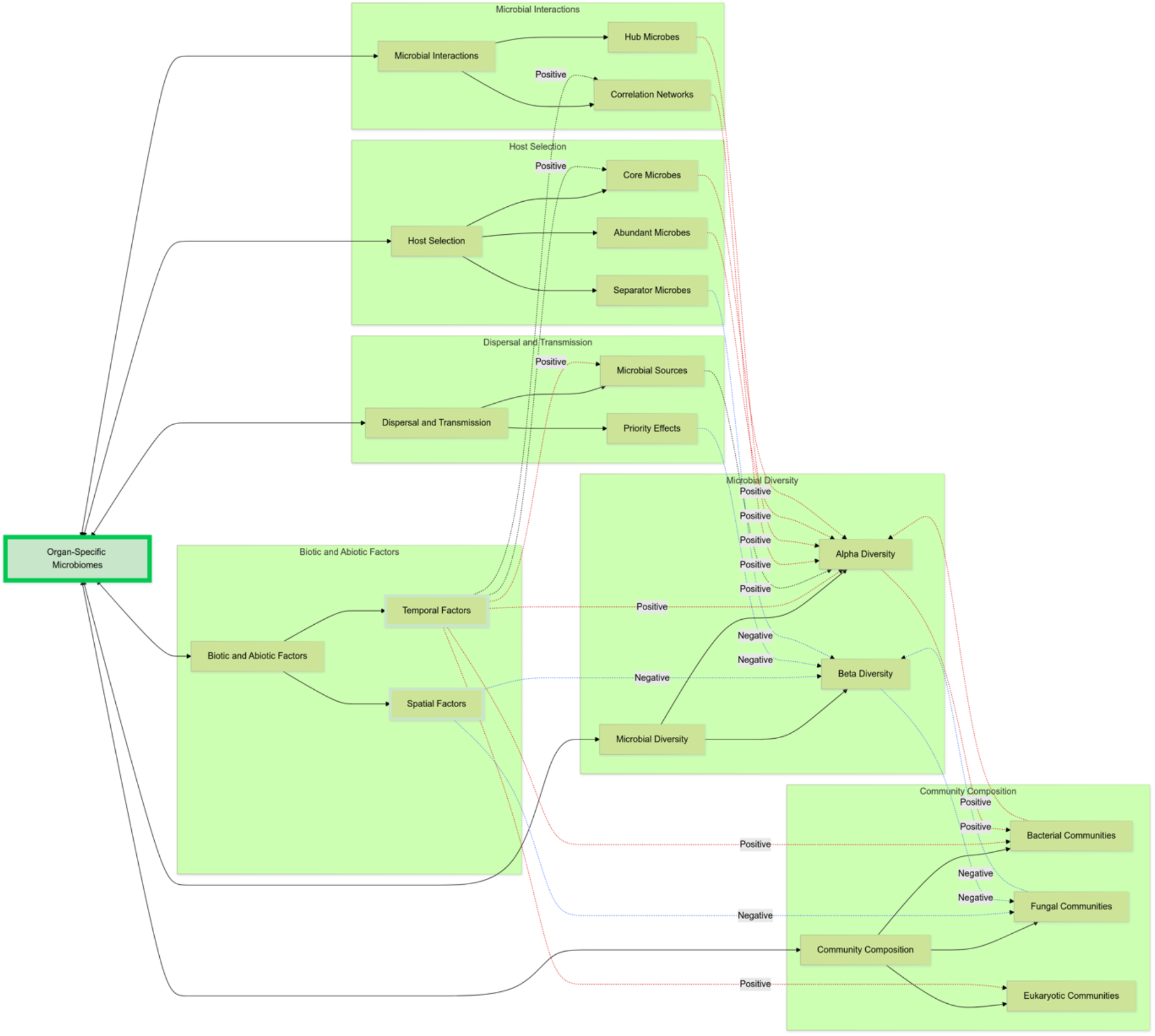
Ecological processes at different tempo-spatial scales that shape metacommunity dynamics in organ-specific *Lotus corniculatus* microbial communities. This figure integrates results from multiyear, multisite sequencing of bacterial, fungal, and eukaryotic communities in *L. corniculatus*, summarizing how spatial, temporal, and ecological processes shape organ-specific microbiomes within a metacommunity framework. Arrows indicate effects supported by results in the manuscript. Solid lines represent hierarchical relationships in the framework (*e.g*., microbial diversity as a component of organ-specificity), while dotted lines indicate functional effects extracted from observed data: red for positive influences and blue for negative influences. For instance, temporal factors positively influenced alpha diversity in aboveground plant parts, whereas spatial factors had a stronger effect on the beta diversity of fungal and eukaryotic communities. Transmission processes such as microbial sourcing and priority effects further shaped organ-specific microbiomes by introducing stochastic variation and assembly history-dependent patterns.

*L. corniculatus* organs host distinct but overlapping microbial communities linked via transmission of microorganisms within the plant host and the outside environment, which signifies that the organs are discrete ecological niches that are interconnected with each other and with the environment. *L. corniculatus* organ-associated microorganisms can also potentially influence community composition during dispersal via priority effects. Consistent with previous observations, the root microbiomes are the most diverse communities among the plant organ communities and have distinct but overlapping community composition with soil microbiomes (42, 99-101). The overlapping and less diverse aboveground microbiomes that are distinct from root and soil microbiomes demonstrated the compositionally-nested characteristic of microbial communities observed in several plant species, where the aboveground communities are subsets of the more diverse belowground communities (9, 99, 100). While the main source of microbiome variation is plant compartment, biotic and abiotic factors from the environment also contribute to patterns of community structures in *L. corniculatus* microbiomes. Plant microbial community structure is also shaped by environmental gradients such as local site conditions, land use, and soil properties, as well as biotic elements like pollinators, insects, and local fauna (10, 97, 102-104). Abiotic and biotic elements in the environment acting at multiple temporal and spatial scales affect *L. corniculatus* microbiomes - while different sampling times and locations affected plant microbial community composition, environmental factors acting at different plant compartments also attributed to the more diverse and stable root microbial communities that are distinct from the less diverse and more variable but overlapping endophytic communities in shoots, flowers, and seeds.

In this study we present bases for future experimentation to explore mechanisms on how key members of *L. corniculatus* organ microbiomes influence community dynamics and species interactions. Variations in community diversity and composition observed at tempo-spatial scales (*i.e*. from differences between root and aboveground plant microbiomes to between year/site variations) provided basis for more in-depth investigation of the crucial roles of abiotic environmental factors, such as soil properties like land use history and soil chemistry, climate and temperature differences in soil and aboveground, or elevation gradients, in shaping these community patterns. This study also provided a requisite basis to test predictions on important agents of horizontal transmission in the environment such as pollinators or on seed microbiomes as initial colonizers during vertical transmission. While in this study we focused on selective filtering by plant organs, microbial interactions, and environmental factors, as well as on stochastic transmission of microorganisms from various sources, other factors such as plant genotype diversity, wider range of geographic locations, or other ecological processes such as genetic drift and diversification of microbiome members additionally cause variations in community dynamics of plant-associated microbiomes. Given the functions of plant-associated microbial communities in plant growth, stress tolerance, and protection, reconstruction of plant microbiomes offers prospects to maximize their beneficial effects for plant productivity, resilience, and pathogen defense. To successfully control plant microbiomes in the field, there remains a need for a comprehensive knowledge of the ecological processes and microbial interactions that shape microbial community dynamics and assembly in natural environments.

## ACKNOWLEDGEMENTS

We thank Bossdorf Lab and Kemen Lab for participating in the sampling trips, especially Elke Klenk for helping in collecting and processing the samples. Through a cooperation agreement with Biodiversity Exploratories (DFG Priority Program 1374) we were granted access to the Schwäbische Alb plots. We thank Ralf Lauterbach and Jörg Hailer from the Schwäbische Alb local management team of the Biodiversity Exploratories for instructions and support during the field work. We thank the managers of the Schwäbische Alb Exploratory, Kirsten Reichel-Jung and Julia Bass and all former managers for their work in maintaining the plot and project infrastructure, Victoria Grießmeier for giving support through the central office, Andreas Ostrowski for managing the central database, and Markus Fischer, Eduard Linsenmair, Dominik Hessenmöller, Daniel Prati, Ingo Schöning, François Buscot, Ernst-Detlef Schulze, Wolfgang W. Weisser and the late Elisabeth Kalko for their role in setting up the Biodiversity Exploratories project. We thank the administration of the UNESCO Biosphere Reserve Swabian Alb as well as all land owners for the excellent collaboration. Field work permits were issued by the responsible state environmental office of Baden-Württemberg.

## AUTHOR CONTRIBUTIONS

E.K., O.B., J.A., and K.L. conceptualized this work. K.L., F.R., and J.A. collected and processed samples from the field. K.L. performed the experiments and data analysis, with M.M. in the machine-learning analysis. K.L., M.M., and E. K. wrote the manuscript, with contributions from all authors. All authors read and approved the final manuscript.

