## Supplementary Figures, Tables, and Methods for "Organ-specific microbiomes in natural *Lotus corniculatus* populations: Metacommunity dynamics in the plant endosphere"

**Supplementary Figure 1.** Increase of target **(a)** bacterial 16S rRNA, **(b)** fungal ITS2, and **(c)** eukaryotic 18S rRNA reads and decrease of corresponding reads of nontarget host *Lotus corniculatus* chloroplast, mitochondria, plant ITS2, and plant 18S rRNA when blocking oligos are used during library preparation of test sequencing samples.

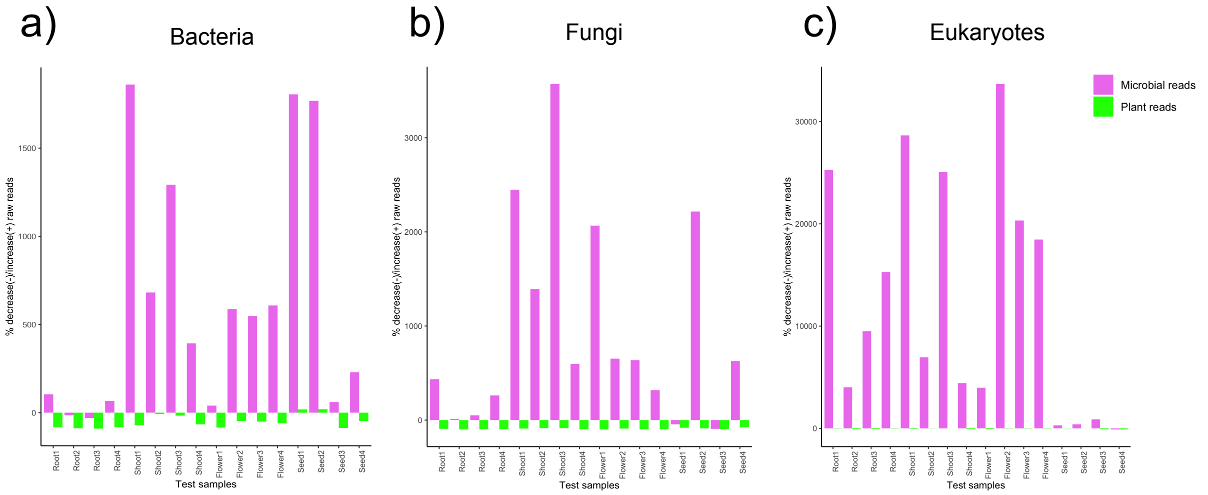


**Supplementary Figure 2**. Boxplots of Observed OTUs of **(a)** bacterial, **(b)** fungal, and **(c)** eukaryotic microbial communities associated with soil and *Lotus corniculatus* plant organs. Datasets have non-normal distribution based on Shapiro-Wilk normality tests ($p<0.05$) thus Kruskal-Wallis rank sum tests were used to test significant differences of $\alpha$-diversity measurements between soil and plant organ samples. Compartments are significantly different if letters are not similar based on post-hoc analysis via Dunn's (Supplementary Table 4).

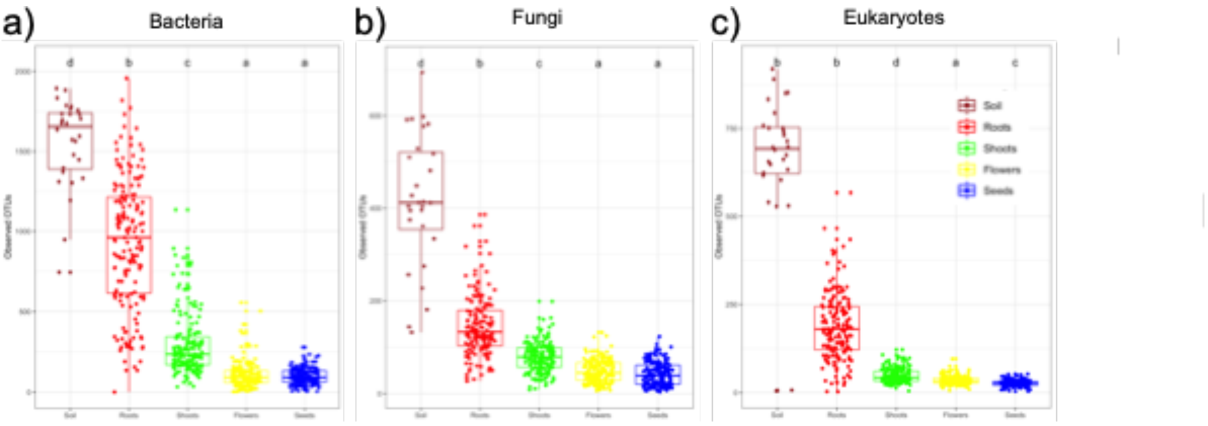

**Supplementary Figure 3.** Relative abundance of the top ten most abundant bacterial, fungal, and eukaryotic phyla detected in all soil and plant samples collected from **(a)** seven grassland sites for **(b)** four years. The relative abundance of the OTUs for each sample were aggregated at the phylum level.

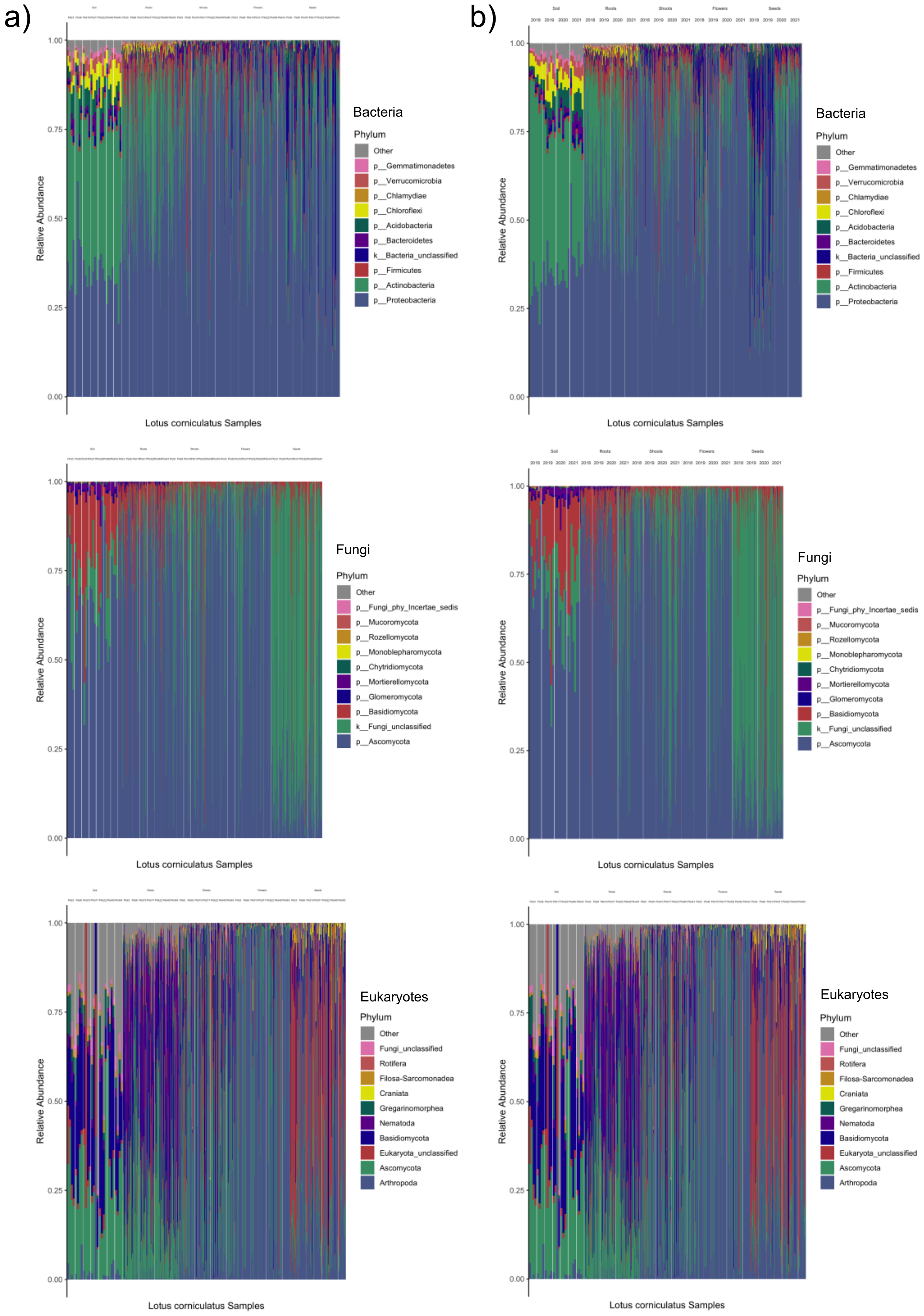

**Supplementary Figure 4.** Barplots of $R^2$ statistic from PERMANOVA showing the percentage of variance that can be explained by factors such as plant organ, year of collection, and sampling sites (Supplementary Table 4).

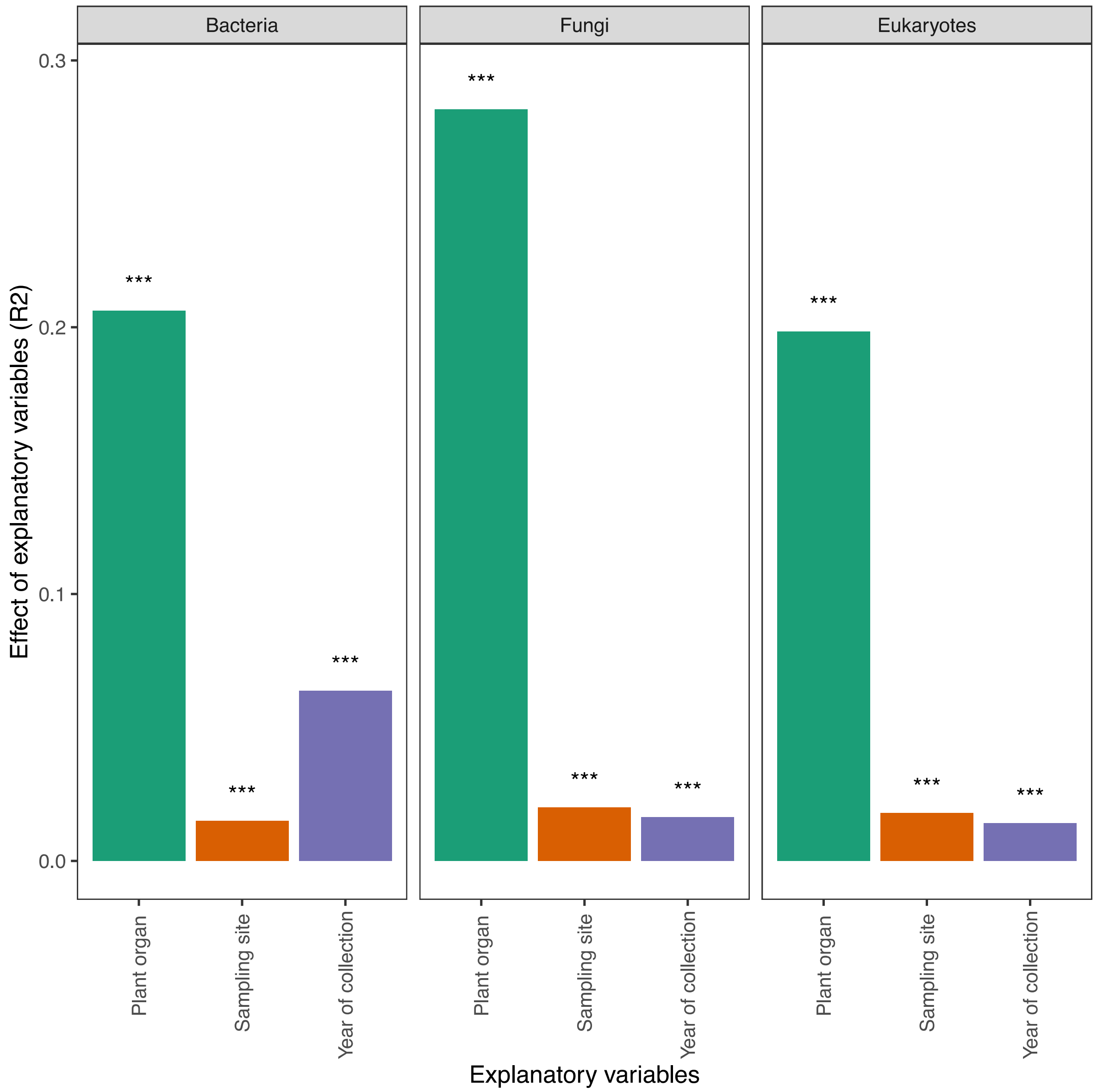

**Supplementary Figure 5.** Discriminatory OTUs that distinguish each organ from others. The OTUs were identified using an SVM classifier with recursive feature elimination and cross-validation. A total of 84, 102, 57, and 166 OTUs for roots, shoots, flowers, and seeds, respectively, were identified. The x-axis values represent the coefficient value of each OTU in separating a specific organ (values < 0) from the rest (values > 0). Only OTUs with absolute coefficient values greater than 0.5 are included.

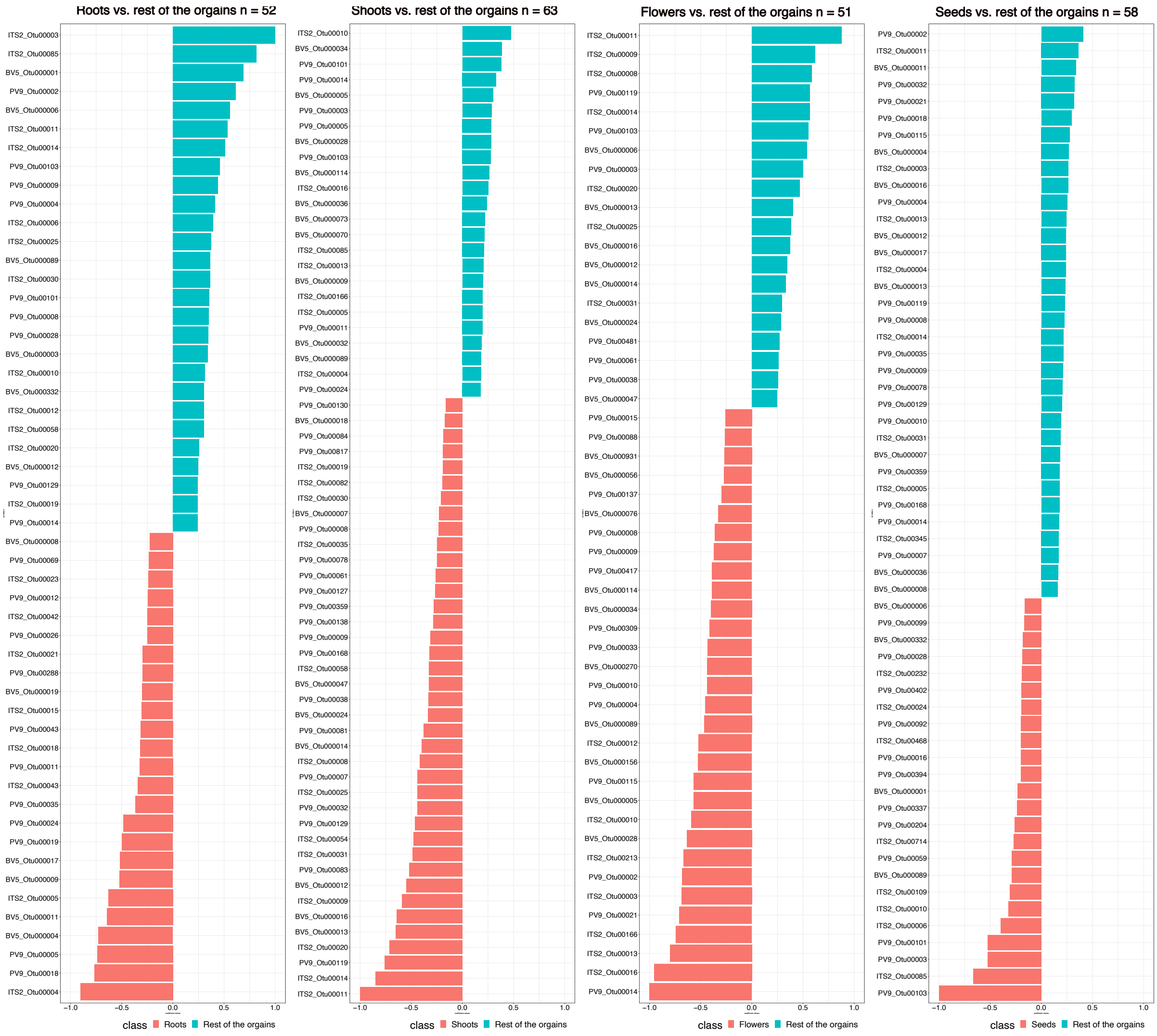

**Supplementary Figure 6**. Microbial hubs of **(a)** root, **(b)** shoot, **(c)** flower, and **(d)** seed microbiomes. Bacterial, fungal, and eukaryotic hubs were identified based on their high betweenness centrality and closeness centrality in correlation networks calculated with FastSpar (Fig. 3). Dotted lines mark the OTUs that are top 5% in betweenness centrality and closeness centrality scores and thus assigned as hub microbes. Size of the circles represent the OTU abundance and green circles are the core microbes.

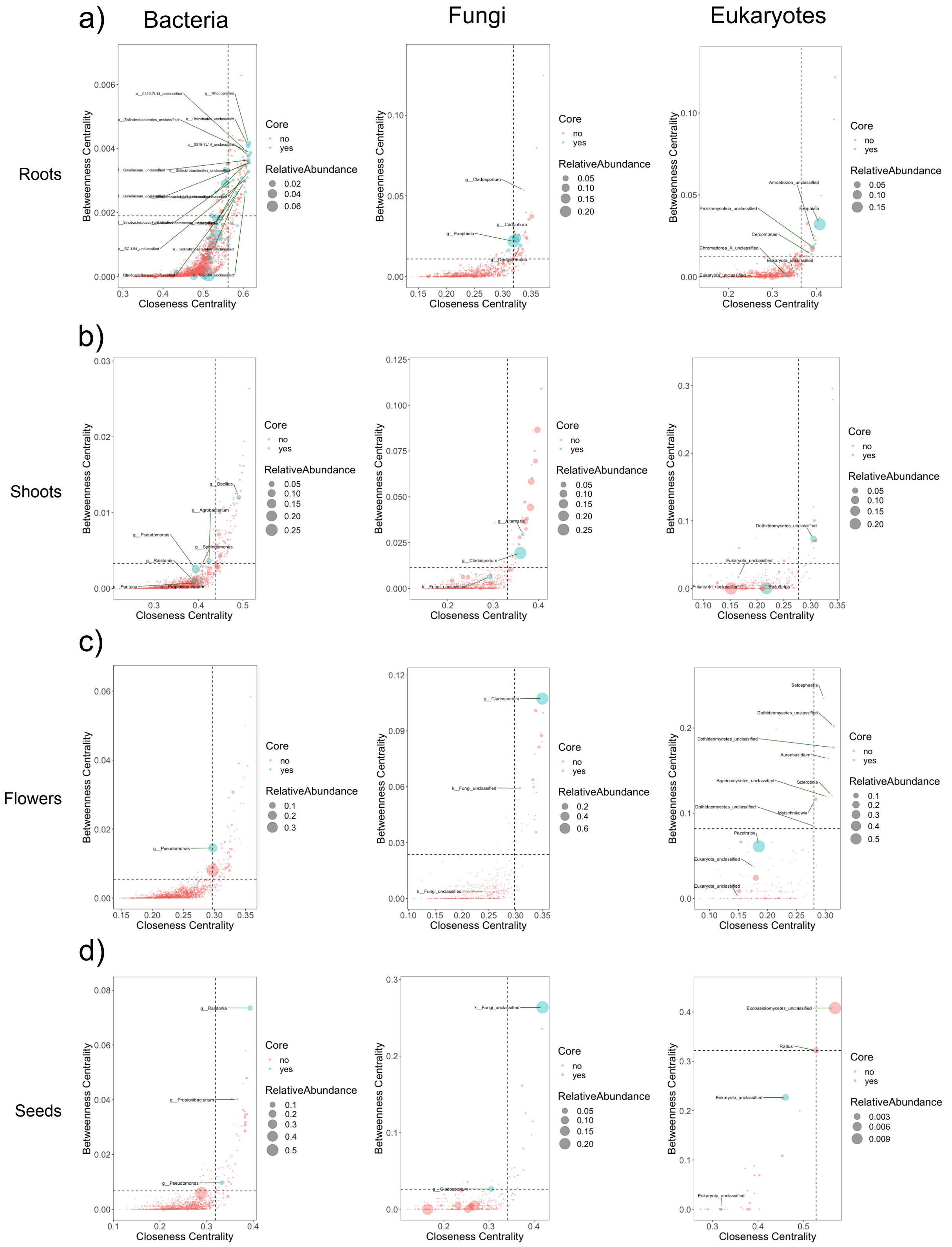

**Supplementary Figure 7.** Boxplots of Shannon's $\alpha$-diversity measurements of bacterial, fungal, and eukaryotic microbial communities associated with whole plants. Shapiro-Wilk normality and Kruskal-Wallis rank sum tests were used to test significant differences of Shannon measurements between **(a-c)** sampling sites and (**d-f)** sampling years. Post-hoc analysis via Dunn's tests indicate that groups are significantly different if letters are not similar.

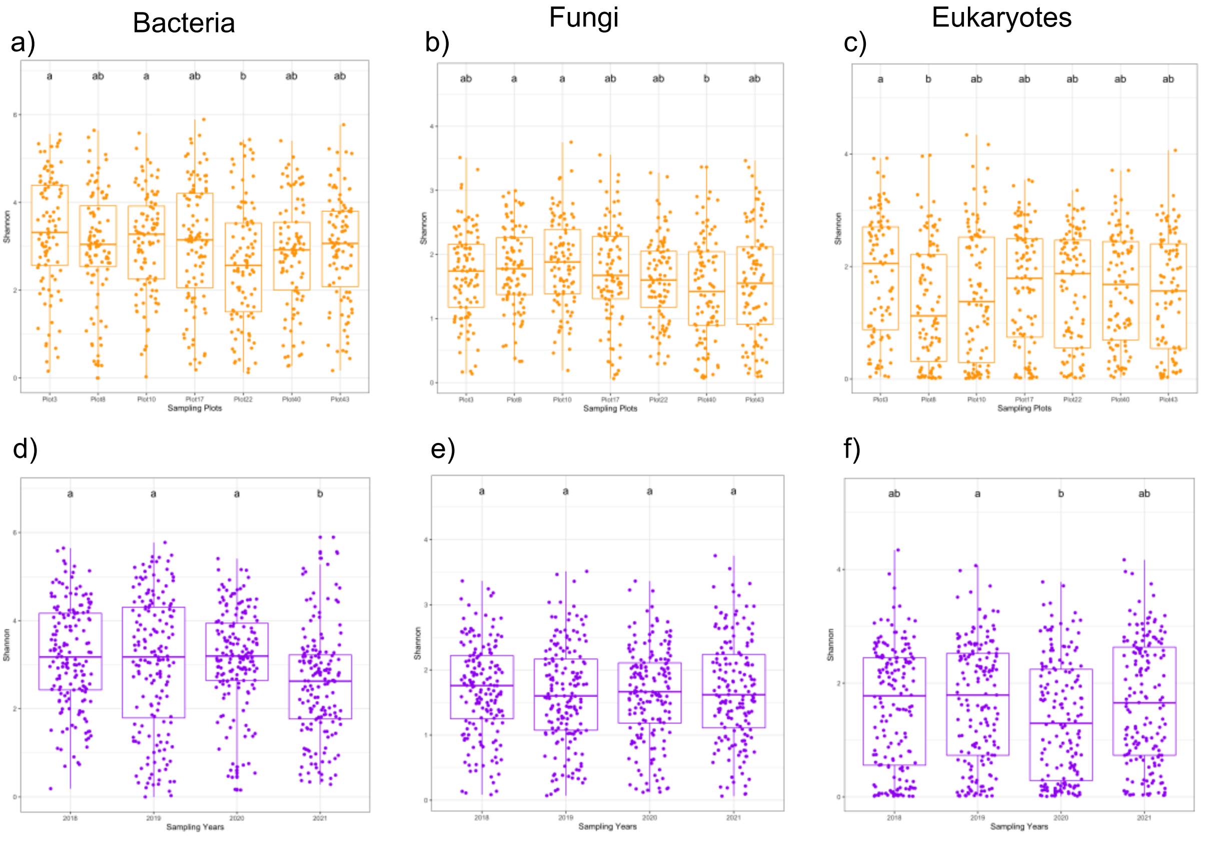

**Supplementary Figure 8.** Relative abundance profiles showing the top most abundant bacterial, fungal, and eukaryotic phyla in plant samples collected from **(a)** seven grassland sites for **(b)** four years. The relative abundance of the OTUs for each site or year were aggregated at the phylum level.

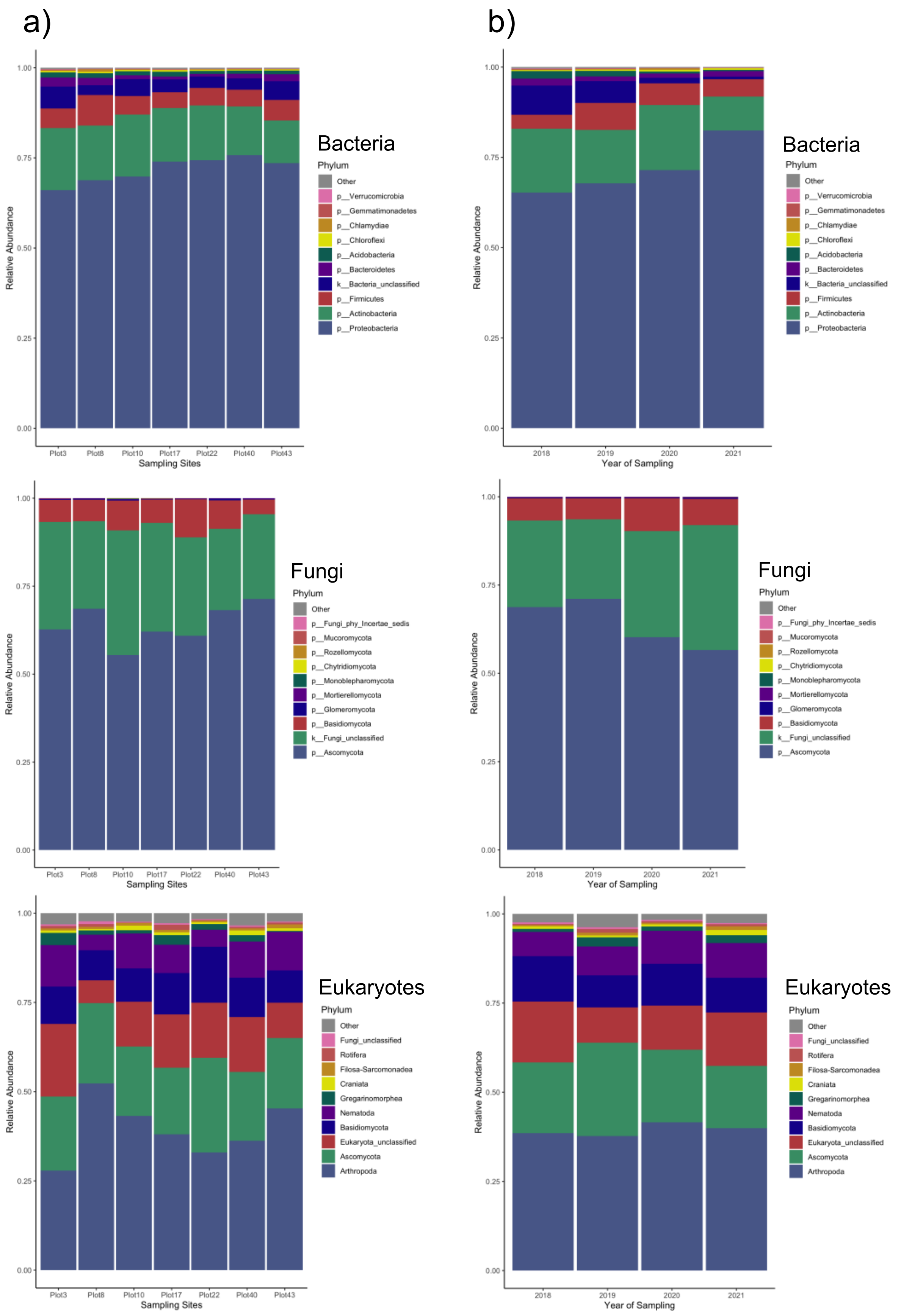

**Supplementary Figure 9.** Principal coordinate plots based on Bray-Curtis dissimilarities between **(a)** bacterial, **(b)** fungal, and **(c)** eukaryotic microbial communities associated with soil and plant organ samples.

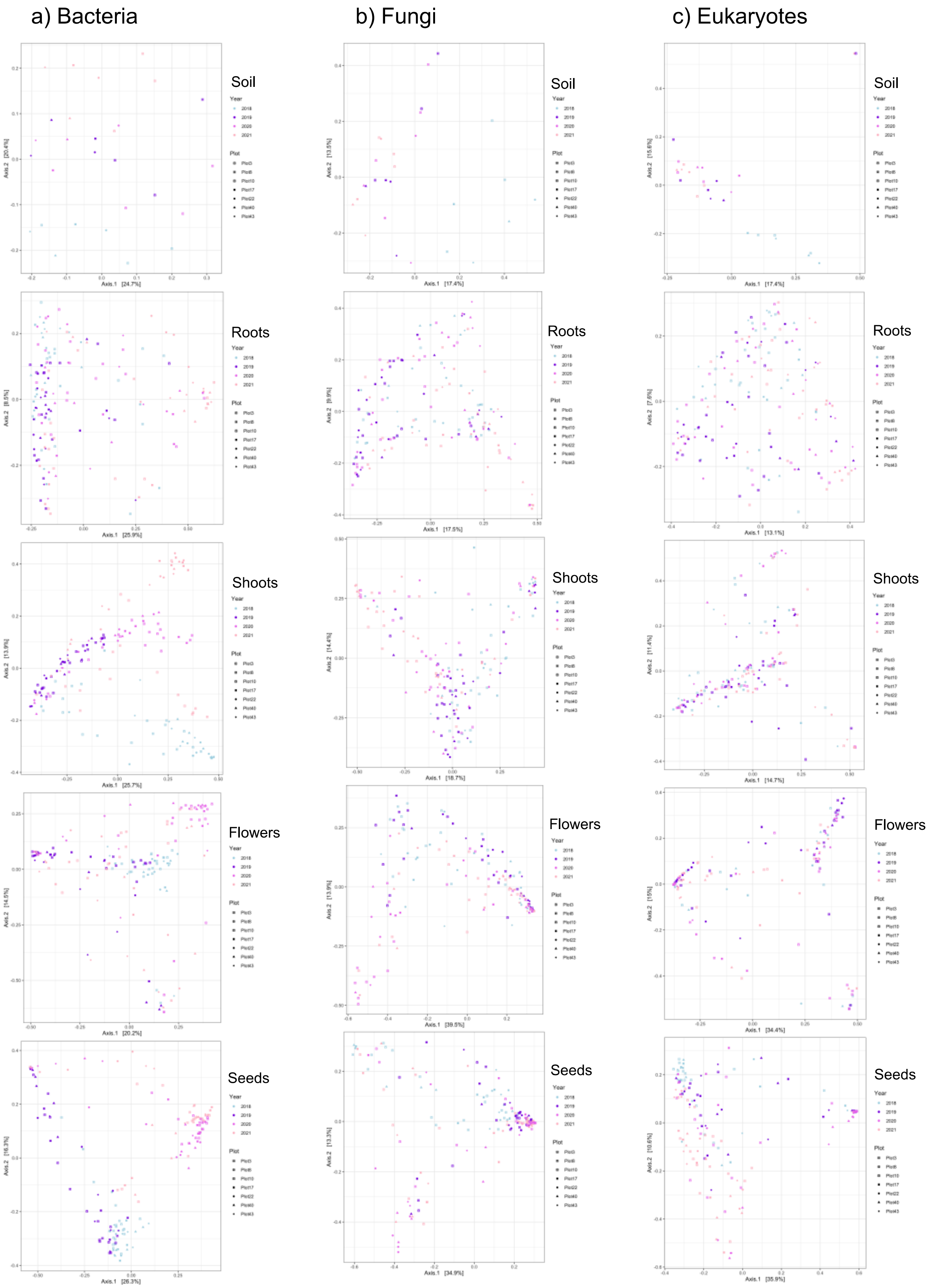

**Supplementary Figure 10.** Distribution of soil and plant **(a)** bacterial, **(b)** fungal, and **(c)** eukaryotic OTUs in soil and plant organs. OTUs are designated as soil OTUs if they are found in soil samples, and the rest of non-soil OTUs are designated as plant OTUs.

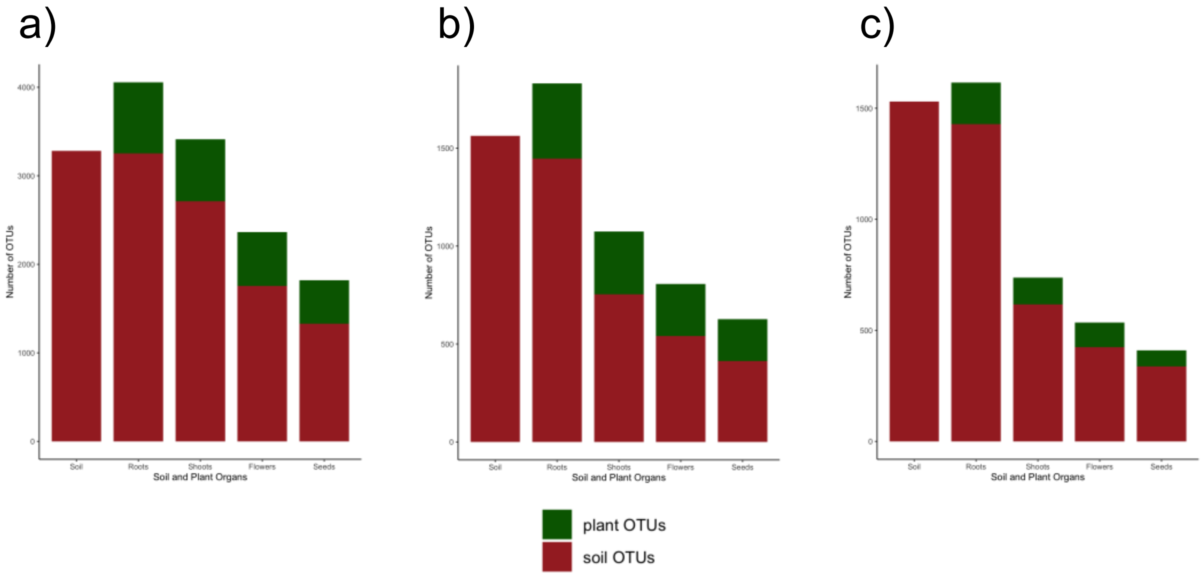


**Supplementary Figure 11**. FEAST calculations of source contribution to the sink microbiomes. Microbial transmission was tracked at multiple directions of sources and sinks. The contribution of the source to the sink microbiomes are in percentages of the total bacterial, fungal, and eukaryotic sink microbiomes, respectively.

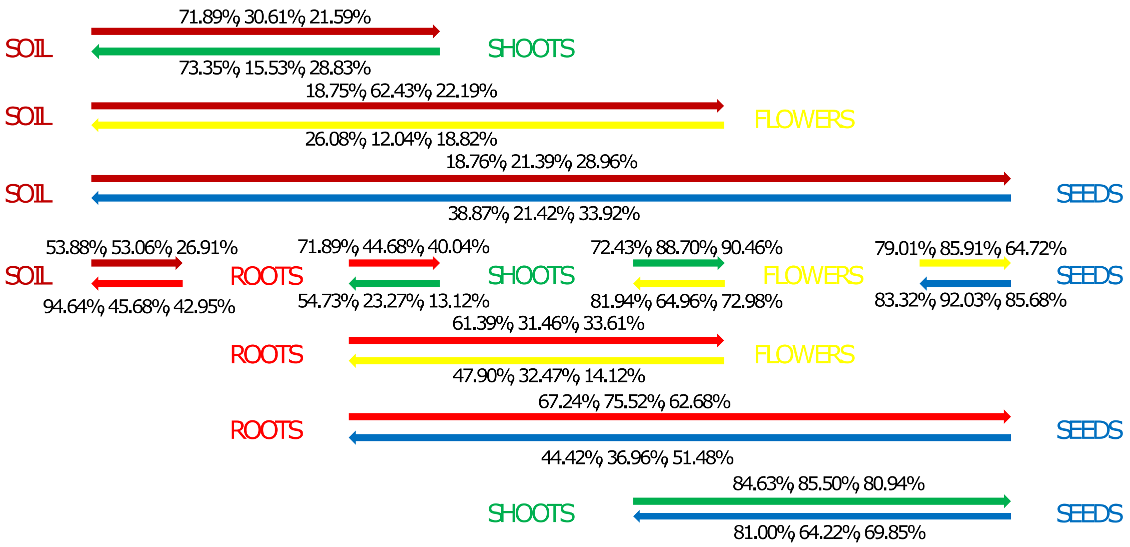

**Supplementary Figure 12**. **(a)** PCoA of Bray-Curtis dissimilarities of root microbial communities and **(b)** PERMANOVA using abundance of candidate early-arriving OTU as explanatory variable of community composition variation. **(a)** and **(b)** show that candidate early-arriving OTUs (BV5_OTU4_*Phyllobacteriaceae,* BV5_OTU17_*Mesorhizobium*) potentially altered the microbial community composition in the roots based on their abundance. (Explanatory variables: abundant OTU: Relative abundance in the root microbial community (RA) >= 0.01; intermediate OTU: RA >= 0.001 and < 0.01; rare OTU: RA < 0.001).

a)

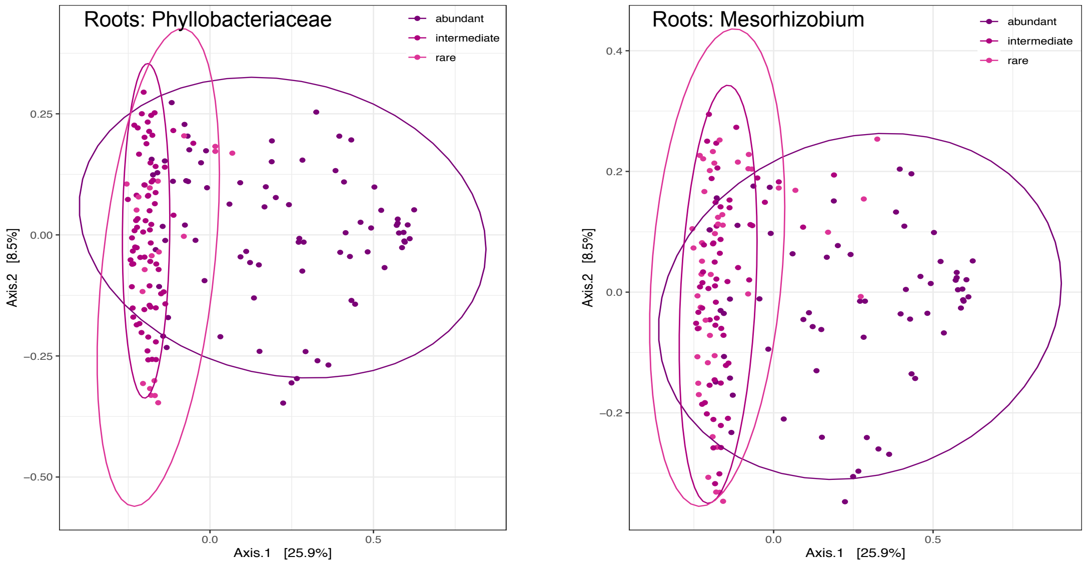


b)

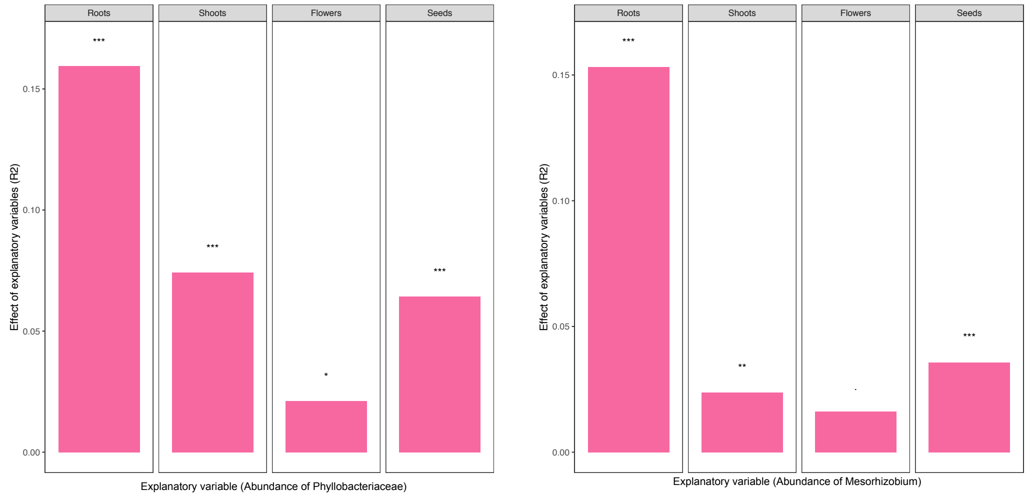

SUPPLEMENTARY TABLES

**Supplementary Table 1.** Primers and blocking oligos used in this study.

| Primer name | Primer sequence (5'-to-3' orientation) |
|---|---|
| 799F | AACMGGATTAGATACCCKG |
| 1192R | ACGTCATCCCCACCTTCC |
| fITS7 | GTGARTCATCGAATCTTTG |
| ITS4 | TCCTCCGCTTATTGATATGC |
| F1422 | ATAACAGGTCTGTGATGCCC |
| R1797 | TGATCCTTCTGCAGGTTCACCTAC |
| clamp1_BV5_mitoF | GATGAGTGTTCGCCCTTGGTCTACGTGGAT |
| clamp1_BV5_mitoR | CTGCTCAGGGTTCCAAACTCAACGTTGGCA |
| clamp1_ITS2_F | AACCATTAGGTCGAGGGCACGTCTGCCTGG |
| clamp1_ITS2_R | TGAGMGYGGTTACACCACGCATGCGGGTCT |
| clamp9_PV9_F | GATGTATTCAACGAGTCTATAGCCTTGGCC |
| Clamp15_PV9_R | TCTCACAACGTCGCAGGCAGCGAACCGCCC |

**Supplementary Table 2.** Basic statistics of the Illumina sequencing data. Amplicon sequencing of the microbial 16S rRNA, ITS2, and 18S rRNA genes in a total of 700 samples of soil and *Lotus corniculatus* roots, shoots, flowers, and seeds, along with blank samples collected from seven grassland sites in the Swabian Alps, Germany for four years.

| Sequencing runs | Sequence reads | Sequence length | %GC |
|---|---|---|---|
| RunLotus1 | 15,367,467 | 300 | 52 |
| RunLotus2 | 19,049,132 | 300 | 52 |
| RunLotus3 | 19,931,059 | 300 | 52.5 |
| RunLotus4 | 18,326,536 | 300 | 52.5 |
| RunLotus6 | 21,112,668 | 301 | 53 |
| RunLotus7 | 20,429,103 | 301 | 52.5 |
| RunLotus8 | 20,404,891 | 301 | 53 |
| RunLotus9 | 21,088,582 | 301 | 52.5 |
| | 155,709,438 (total) | | |

**Supplementary Table 3.** Summary of sequence data after processing by mothur and phyloseq.

| Locus | Domain | Subdomain | Kingdom | Phylum | Subphylum | Class | Order | Family | Genus | Species | OTUs |
|---|---|---|---|---|---|---|---|---|---|---|---|
| 16S rRNA | - | - | Bacteria | 25 | - | 95 | 182 | 335 | 586 | 671 | 4,225 |
| ITS2 | - | - | Fungi | 15 | - | 51 | 129 | 280 | 531 | 788 | 2,027 |
| 18S rRNA | Eukaryota | 10 | 26 | 73 | 122 | 197 | - | - | 425 | 486 | 1,773 |

**Supplementary Table 4.** Statistical analysis of alpha- and beta-diversity measurements

**(a)** Shapiro-Wilk normality tests

| $\alpha$-diversity index | Statistic (W) | p-value | normality |
|---|---|---|---|
| 16S rRNA_Shannon | 0.98747 | 1.091e-05 | non-normal distribution |
| 16S rRNA_Observed | 0.75878 | < 2.2e-16 | non-normal distribution |
| ITS2_Shannon | 0.98424 | 7.343e-07 | non-normal distribution |
| ITS2_Observed | 0.70022 | < 2.2e-16 | non-normal distribution |
| 18S rRNA_Shannon | 0.95252 | 3.048e-14 | non-normal distribution |
| 18S rRNA_Observed | 0.56119 | < 2.2e-16 | non-normal distribution |

**(b)** Kruskal-Wallis rank sum tests by soil/plant organs and post-hoc analysis via Dunn's test

| $\alpha$-diversity index | Statistic (K-W chi-squared) | p-value | significance |
|---|---|---|---|
| 16S rRNA_Shannon | 295.25 | < 2.2e-16 | significant differences |

| | | | |
|---|---|---|---|
| 16S rRNA_Observed | 498.33 | < 2.2e-16 | significant differences |
| ITS2_Shannon | 237.31 | < 2.2e-16 | significant differences |
| ITS2_Observed | 397.04 | < 2.2e-16 | significant differences |
| 18S rRNA_Shannon | 258.82 | < 2.2e-16 | significant differences |
| 18S rRNA_Observed | 423.6 | < 2.2e-16 | significant differences |

**(c)** Permutational multivariate analysis of variance (PERMANOVA) of variables plant organ, collection year, and sampling sites.
Permutation test for adonis under reduced model
Terms added sequentially (first to last)
Permutation: free
Number of permutations: 999

| 16S rRNA | | | | | |
|---|---|---|---|---|---|
| | Df | SumOfSqs | R2 | F | Pr(>F) |
| PlantOrgan | 4 | 55.372 | 0.20622 | 49.3158 | 0.001 *** |
| Year | 3 | 17.103 | 0.06370 | 20.3100 | 0.001 *** |
| Plot | 6 | 4.029 | 0.01501 | 2.3923 | 0.001 *** |
| Residual | 684 | 192.001 | 0.71507 | | |
| Total | 697 | 268.506 | 1.00000 | | |
| ITS2 | | | | | |
| | Df | SumOfSqs | R2 | F | Pr(>F) |
| PlantOrgan | 4 | 76.128 | 0.28167 | 70.9434 | 0.001 *** |
| Year | 3 | 4.441 | 0.01643 | 5.5178 | 0.001 *** |
| Plot | 6 | 5.408 | 0.02001 | 3.3598 | 0.001 *** |
| Residual | 687 | 184.303 | 0.68190 | | |
| Total | 700 | 270.280 | 1.00000 | | |
| 18S rRNA | | | | | |
| | Df | SumOfSqs | R2 | F | Pr(>F) |
| PlantOrgan | 4 | 60.755 | 0.19845 | 44.2882 | 0.001 *** |
| Year | 3 | 4.310 | 0.01408 | 4.1894 | 0.001 *** |
| Plot | 6 | 5.475 | 0.01788 | 2.6606 | 0.001 *** |
| Residual | 687 | 235.610 | 0.76959 | | |
| Total | 700 | 306.151 | 1.00000 | | |

Significance codes: 0 ‘***’ 0.001 ‘**’ 0.01 ‘*’ 0.05 ‘.’ 0.1 ‘ ’ 1
Abbreviations: Df ‘degrees of freedom’; SumsOfSqs ‘Sums of Squares’; F ‘F statistic’; R2 ‘R2 statistic’; Pr(>F) ‘p-values’.

**Supplementary Table 5.** Performance tests of the SVM models.

| Taxa | Organ | | precision | recall | f1-score | support |
|---|---|---|---|---|---|---|
| Bacteria | Seeds | 0 | 0.625 | 0.80357143 | 0.703125 | 56 |
| Bacteria | Flowers | 1 | 0.69047619 | 0.52727273 | 0.59793814 | 55 |
| Bacteria | Shoots | 2 | 0.81132075 | 0.76785714 | 0.78899083 | 56 |
| Bacteria | Roots | 3 | 0.98148148 | 0.96363636 | 0.97247706 | 55 |
| Bacteria | Soil | 4 | 0.9 | 1 | 0.94736842 | 9 |
| Bacteria | | accuracy | 0.77489177 | 0.77489177 | 0.77489177 | 0.77489177 |
| Bacteria | | macro avg | 0.80165569 | 0.81246753 | 0.80197989 | 231 |
| Bacteria | | weighted avg | 0.78134907 | 0.77489177 | 0.77254389 | 231 |
| Taxa | | | precision | recall | f1-score | support |
| Fungi | Seeds | 0 | 0.7 | 0.89090909 | 0.784 | 55 |
| Fungi | Flowers | 1 | 0.75806452 | 0.83928571 | 0.79661017 | 56 |
| Fungi | Shoots | 2 | 0.9 | 0.64285714 | 0.75 | 56 |
| Fungi | Roots | 3 | 1 | 0.91071429 | 0.95327103 | 56 |

| | | | | | | |
|---|---|---|---|---|---|---|
| Fungi | Soil | 4 | 1 | 1 | 1 | 9 |
| Fungi | | accuracy | 0.82758621 | 0.82758621 | 0.82758621 | 0.82758621 |
| Fungi | | macro avg | 0.8716129 | 0.85675325 | 0.85677624 | 232 |
| Fungi | | weighted avg | 0.84634316 | 0.82758621 | 0.82807477 | 232 |
| Taxa | Organ | | precision | recall | f1-score | support |
| Eukaryote | Seeds | 0 | 0.85 | 0.92727273 | 0.88695652 | 55 |
| Eukaryote | Flowers | 1 | 0.73846154 | 0.85714286 | 0.79338843 | 56 |
| Eukaryote | Shoots | 2 | 0.76595745 | 0.64285714 | 0.69902913 | 56 |
| Eukaryote | Roots | 3 | 0.98076923 | 0.91071429 | 0.94444444 | 56 |
| Eukaryote | Soil | 4 | 0.875 | 0.77777778 | 0.82352941 | 9 |
| Eukaryote | | accuracy | 0.83189655 | 0.83189655 | 0.83189655 | 0.83189655 |
| Eukaryote | | macro avg | 0.84203764 | 0.82315296 | 0.82946959 | 232 |
| Eukaryote | | weighted avg | 0.8353256 | 0.83189655 | 0.8304252 | 232 |

**Supplementary Table 6.** Abundant OTUs (**relative abundance > 1%**) in *Lotus corniculatus* organs.

| | | Relative abundance | | | |
|---|---|---|---|---|---|
| OTU | Genus | Roots | Shoots | Flowers | Seeds |
| Bacteria | | | | | |
| BV5_Otu000001 | *Pantoea* | **0.01498769** | **0.25475602** | **0.39135219** | **0.53456217** |
| BV5_Otu000003 | *Pseudomonas* | **0.01419121** | **0.10772585** | **0.20539426** | **0.0531343** |
| BV5_Otu000004 | *Phyllobacteriaceae* | **0.07248534** | 0.00054067 | 0.00086442 | 0.00054539 |
| BV5_Otu000005 | *Enterobacteriaceae* | 2.45E-05 | 0.00237893 | **0.12226135** | **0.01422907** |
| BV5_Otu000006 | *Ralstonia* | 0.0005517 | **0.06358809** | **0.01562879** | **0.07815603** |
| BV5_Otu000007 | *Bacillus* | **0.03395148** | **0.02838181** | 0.00241461 | 0.00069461 |
| BV5_Otu000008 | *Frankia* | **0.04906623** | 0.00023149 | 3.93E-05 | 1.57E-05 |
| BV5_Otu000009 | *Phyllobacterium* | **0.03919923** | 0.00068449 | 0.00030655 | 7.77E-05 |
| BV5_Otu000010 | *Pantoea* | 0.00091647 | **0.0124075** | **0.01680126** | **0.02267523** |
| BV5_Otu000011 | *Mesorhizobium* | **0.02558929** | 0.00063515 | 0.00072956 | 7.94E-05 |
| BV5_Otu000012 | *Burkholderia* | 0.00164728 | **0.03042322** | 0.00972065 | **0.01644205** |
| BV5_Otu000013 | *Burkholderia* | 0.00083817 | **0.03141899** | 0.00876367 | **0.01736878** |
| BV5_Otu000014 | *Agrobacterium* | 0.00501434 | **0.03865919** | 0.00136009 | 0.00281248 |
| BV5_Otu000015 | *Streptomyces* | **0.02373183** | 0.00045353 | 9.95E-05 | 0.00011431 |
| BV5_Otu000016 | *Burkholderia* | 0.00074167 | **0.0303996** | 0.00767331 | **0.01618725** |
| BV5_Otu000017 | *Mesorhizobium* | **0.02286487** | 0.00031075 | 0.00065989 | 0.00025132 |
| BV5_Otu000018 | *Comamonadaceae* | 0.0062342 | **0.01769185** | 0.00033791 | **0.02457407** |
| BV5_Otu000019 | *Cryptosporangium* | **0.02155104** | 0.00015905 | 0.00010301 | 2.01E-05 |
| BV5_Otu000021 | *Pantoea* | 0.00052872 | 0.00695622 | **0.01308926** | **0.01592197** |
| BV5_Otu000022 | *Methylibium* | **0.019637** | 0.00022047 | 7.02E-05 | 1.83E-05 |
| BV5_Otu000023 | *Steroidobacter* | **0.01899183** | 0.00025301 | 5.92E-05 | 4.80E-05 |
| BV5_Otu000024 | *Xanthomonadaceae* | 0.00477739 | **0.02151746** | 0.00025778 | 0.00059513 |
| BV5_Otu000028 | *Wolbachia* | 1.01E-06 | 0.0014283 | **0.01519235** | **0.01760352** |
| BV5_Otu000032 | *Pseudomonas* | 0.00045571 | 0.00352115 | **0.01337044** | 0.00700109 |
| BV5_Otu000040 | *Sodalis* | 0.00184406 | 1.26E-05 | 5.47E-06 | **0.02810647** |
| BV5_Otu000056 | *Enterobacteriaceae* | 0.00108648 | 0.00033332 | **0.01385366** | 3.49E-05 |
| BV5_Otu000068 | *Gluconobacter* | 2.27E-06 | 5.25E-06 | **0.0128056** | 1.75E-06 |
| Fungi | | | | | |
| ITS2_Otu00003 | *Cladosporium* | 0.00108977 | **0.2671789** | **0.76240753** | **0.04540433** |
| ITS2_Otu00004 | *Exophiala* | **0.23861407** | 0.00042552 | 0.00037056 | 0.00037517 |
| ITS2_Otu00005 | *Cadophora* | **0.12629519** | 0.00027117 | 0.0001305 | 0.00026054 |
| ITS2_Otu00006 | *Fungi* | **0.01084811** | **0.04378309** | **0.01783285** | **0.24165332** |
| ITS2_Otu00008 | *Boeremia* | 0.00022675 | **0.06855963** | 0.00358989 | **0.15808927** |
| ITS2_Otu00009 | *Didymellaceae* | 0.00791914 | **0.0633385** | **0.01207996** | 0.00837013 |
| ITS2_Otu00010 | *Sclerotiniaceae* | 1.36E-05 | 2.38E-05 | **0.03687987** | **0.20319659** |

| | | | | | |
|---|---|---|---|---|---|
| ITS2_Otu00011 | *Septoria* | 4.2685E-05 | **0.09075922** | 5.94E-05 | 0.0002119 |
| ITS2_Otu00012 | *Alternaria* | 4.83E-05 | **0.02015062** | **0.03302806** | **0.0907913** |
| ITS2_Otu00013 | *Didymellaceae* | 0.00023233 | **0.02319715** | **0.03025786** | **0.03560474** |
| ITS2_Otu00014 | *Chaetosphaeronema* | 0.00059344 | **0.07000393** | 4.40E-05 | 0.00016153 |
| ITS2_Otu00015 | *Pseudomassaria* | **0.02789404** | 4.14E-05 | 3.9402E-05 | 2.95E-05 |
| ITS2_Otu00016 | *Botryotinia* | 2.97E-05 | 0.00139567 | **0.04266852** | 0.00042554 |
| ITS2_Otu00017 | *Helotiales* | **0.02200804** | 0.00236582 | 1.64E-05 | 9.90E-05 |
| ITS2_Otu00018 | *Tetracladium* | **0.02145356** | 1.11E-05 | 1.59E-05 | 1.74E-05 |
| ITS2_Otu00019 | *Alternaria* | 4.93E-05 | **0.01773128** | **0.01641507** | **0.03986536** |
| ITS2_Otu00020 | *Podospora* | 1.12E-05 | **0.04191523** | 0.00152255 | 3.47E-05 |
| ITS2_Otu00021 | *Dactylonectria* | **0.01881724** | 1.4032E-05 | 1.59E-05 | 4.86E-05 |
| ITS2_Otu00022 | *Neocosmospora* | **0.0187555** | 2.68E-05 | 4.10E-06 | 1.39E-05 |
| ITS2_Otu00023 | *Cistella* | **0.01668841** | 8.16E-06 | 5.79E-05 | 3.65E-05 |
| ITS2_Otu00024 | *Gibberella* | 0.00182659 | **0.01768527** | 0.00076435 | **0.07204492** |
| ITS2_Otu00025 | *Colletotrichum* | 0.00125693 | **0.03415043** | 0.00052657 | 0.00015979 |
| ITS2_Otu00026 | *Mycena* | **0.0148273** | 6.79E-05 | 1.25E-05 | 1.74E-05 |
| ITS2_Otu00028 | *Ascomycota* | **0.01283799** | 0.00064285 | 0.00170248 | 5.21E-05 |
| ITS2_Otu00030 | *Uromyces* | 1.29E-06 | **0.03014028** | 0.00010317 | 2.78E-05 |
| ITS2_Otu00031 | *Stemphylium* | 0.00164826 | **0.02476839** | 4.30E-05 | 0.00101609 |
| ITS2_Otu00085 | *Fungi* | 0.00087376 | 0.00188744 | 0.00146493 | **0.01086953** |
| ITS2_Otu00109 | *Malassezia* | 8.0501E-05 | 0.0006931 | 0.0005539 | **0.02153931** |
| Eukaryotes | | | | | |
| PV9_Otu00002 | *Pezothrips* | 0.00127649 | **0.22407626** | **0.5935281** | 0.00544734 |
| PV9_Otu00003 | *Insecta* | 0.0001378 | 0.00840221 | 0.00466225 | **0.9106856** |
| PV9_Otu00004 | *Frankliniella* | 0.00015456 | **0.09482595** | **0.12677873** | 0.00044784 |
| PV9_Otu00005 | *Exophiala* | **0.18068931** | 0.00018327 | 6.80E-05 | 9.03E-05 |
| PV9_Otu00007 | *Pucciniomycetes* | 0.00063188 | **0.23242558** | 0.0020743 | 0.00016626 |
| PV9_Otu00008 | *Mayetiola* | 0.00012911 | **0.07492251** | **0.0488326** | 0.00012514 |
| PV9_Otu00009 | *Dothideomycetes* | **0.02789218** | **0.06098462** | **0.01528368** | 0.00410025 |
| PV9_Otu00010 | *Aeolothrips* | 2.76E-05 | 0.00940713 | **0.05010144** | 3.66E-05 |
| PV9_Otu00011 | *Agaricomycetes* | **0.05681411** | 6.27E-05 | 2.94E-05 | 3.22E-05 |
| PV9_Otu00012 | *Agaricomycetes* | **0.04652027** | 0.00029151 | 1.51E-05 | 5.99E-05 |
| PV9_Otu00013 | *Insecta* | 1.18E-05 | **0.03836716** | **0.02424527** | 0.00065611 |
| PV9_Otu00014 | *Metschnikowia* | 1.24E-05 | 0.00027306 | **0.0343287** | 5.10E-05 |
| PV9_Otu00015 | *Insecta* | 1.64E-05 | **0.01191882** | **0.02390365** | **0.02006059** |
| PV9_Otu00016 | *Agaricomycetes* | **0.03859289** | 0.0001064 | 5.70E-06 | 0.00031286 |
| PV9_Otu00018 | *Aglenchus* | **0.03959254** | 3.69E-05 | 1.12E-05 | 0.00012067 |
| PV9_Otu00019 | *Agaricomycetes* | **0.02896228** | 0.00206027 | 1.56E-05 | 1.07E-05 |
| PV9_Otu00021 | *Insecta* | 1.64E-05 | 0.00815929 | **0.01215686** | 1.79E-05 |
| PV9_Otu00022 | *Agaricomycetes* | **0.03146715** | 9.84E-06 | 4.61E-06 | 4.92E-05 |
| PV9_Otu00024 | *Chromadorea* | **0.02882231** | 5.66E-05 | 1.51E-05 | 2.41E-05 |
| PV9_Otu00026 | *Pezizomycotina* | **0.02743875** | 1.23E-05 | 8.78E-06 | 2.32E-05 |
| PV9_Otu00032 | *Arachnida* | 4.03E-06 | **0.04404551** | 0.00080172 | 1.34E-05 |
| PV9_Otu00033 | *Arachnida* | 3.26E-05 | 7.38E-06 | **0.01642066** | 1.79E-05 |
| PV9_Otu00038 | *Insecta* | 6.52E-06 | **0.0440246** | 1.54E-06 | 2.86E-05 |
| PV9_Otu00081 | *Leotiomycetes* | 3.57E-05 | **0.01225707** | 0.00010707 | 9.83E-06 |
| PV9_Otu00101 | *Exobasidiomycetes* | 0.00018094 | 0.00033456 | 0.00011497 | **0.01129961** |

**Supplementary Table 7.** Key microbes (abundant OTUs: relative abundance > 1%; core OTUs: persistent in 90% of samples; hub OTUs; separator OTUs: predicted by SVM model) in *Lotus corniculatus* organs.

**(a)** Roots

| | Separator | Abundant | Core | Hub |
|---|---|---|---|---|
| BV5_Otu000001_Pantoea | yes | yes | no | no |
| BV5_Otu000003_Pseudomonas | yes | yes | yes | no |
| BV5_Otu000004_Phyllobacteriaceae | yes | yes | yes | no |
| BV5_Otu000005_Enterobacteriaceae | no | no | no | no |
| BV5_Otu000006_Ralstonia | yes | no | no | no |

| | | | | |
|---|---|---|---|---|
| BV5_Otu000007_Bacillus | no | yes | yes | no |
| BV5_Otu000008_Frankia | yes | yes | yes | no |
| BV5_Otu000009_Phyllobacterium | yes | yes | yes | no |
| BV5_Otu000010_Pantoea | no | no | no | no |
| BV5_Otu000011_Mesorhizobium | yes | yes | yes | no |
| BV5_Otu000012_Burkholderia | yes | no | no | no |
| BV5_Otu000013_Burkholderia | yes | no | no | no |
| BV5_Otu000014_Agrobacterium | no | no | no | no |
| BV5_Otu000015_Streptomyces | no | yes | yes | no |
| BV5_Otu000016_Burkholderia | yes | no | no | no |
| BV5_Otu000017_Mesorhizobium | yes | yes | yes | no |
| BV5_Otu000018_Comamonadaceae | no | no | yes | no |
| BV5_Otu000019_Cryptosporangium | yes | yes | yes | no |
| BV5_Otu000021_Pantoea | no | no | no | no |
| BV5_Otu000022_Methylibium | no | yes | yes | no |
| BV5_Otu000023_Steroidobacter | yes | yes | yes | no |
| BV5_Otu000024_Xanthomonadaceae | no | no | yes | no |
| BV5_Otu000025_Rhodoplanes | no | no | yes | yes |
| BV5_Otu000026_Rhizobium | yes | yes | yes | no |
| BV5_Otu000027_Gaiellaceae | no | no | yes | yes |
| BV5_Otu000028_Wolbachia | no | no | no | no |
| BV5_Otu000029_Rhizobiales | no | no | yes | yes |
| BV5_Otu000030_0319-7L14 | no | no | yes | yes |
| BV5_Otu000032_Pseudomonas | no | no | no | no |
| BV5_Otu000033_Chryseobacterium | no | yes | no | no |
| BV5_Otu000034_Prauserella | no | no | no | no |
| BV5_Otu000036_Bosea | yes | yes | yes | no |
| BV5_Otu000039_Methylibium | no | no | yes | no |
| BV5_Otu000040_Sodalis | no | no | no | no |
| BV5_Otu000043_Bradyrhizobiaceae | no | no | yes | no |
| BV5_Otu000045_Micromonosporaceae | no | no | yes | no |
| BV5_Otu000047_Methylobacterium | no | no | no | no |
| BV5_Otu000049_Kineosporia | no | no | yes | no |
| BV5_Otu000052_Methylobacterium | yes | no | no | no |
| BV5_Otu000054_Gaiellaceae | no | no | yes | yes |
| BV5_Otu000055_Methylobacterium | no | no | no | no |
| BV5_Otu000056_Enterobacteriaceae | no | no | no | no |
| BV5_Otu000057_Rhodospirillaceae | no | no | yes | yes |
| BV5_Otu000059_Solirubrobacterales | no | no | yes | yes |
| BV5_Otu000060_Hyphomicrobium | no | no | yes | no |
| BV5_Otu000061_Mycobacterium | no | no | yes | no |
| BV5_Otu000062_Flavobacterium | no | no | yes | no |
| BV5_Otu000063_SC-I-84 | no | no | yes | yes |
| BV5_Otu000065_Solirubrobacterales | no | no | yes | yes |
| BV5_Otu000068_Gluconobacter | no | no | no | no |
| BV5_Otu000069_Pseudomonas | no | no | no | no |
| BV5_Otu000070_Rubrobacter | no | no | no | no |
| BV5_Otu000073_Erwinia | no | no | no | no |
| BV5_Otu000075_Solirubrobacteraceae | no | no | yes | yes |
| BV5_Otu000076_Acinetobacter | no | no | no | no |
| BV5_Otu000077_Solirubrobacterales | no | no | yes | yes |
| BV5_Otu000078_Sinobacteraceae | no | no | yes | yes |
| BV5_Otu000079_Buchnera | no | no | no | no |
| BV5_Otu000080_Pedomicrobium | no | no | yes | no |
| BV5_Otu000081_SC-I-84 | no | no | yes | yes |
| BV5_Otu000084_Dongia | no | no | yes | no |
| BV5_Otu000086_Burkholderiaceae | no | no | no | no |
| BV5_Otu000089_Propionibacterium | yes | no | no | no |

| | | | | |
|---|---|---|---|---|
| BV5_Otu000091_Patulibacteraceae | no | no | yes | no |
| BV5_Otu000092_0319-7L14 | no | no | yes | yes |
| BV5_Otu000094_Caulobacter | no | no | yes | no |
| BV5_Otu000095_Sphingomonas | no | no | no | no |
| BV5_Otu000096_Bradyrhizobium | no | no | yes | no |
| BV5_Otu000097_Gaiellaceae | no | no | yes | yes |
| BV5_Otu000098_Mycobacterium | no | no | yes | no |
| BV5_Otu000102_Solibacillus | no | no | yes | no |
| BV5_Otu000105_Labrys | no | no | yes | no |
| BV5_Otu000106_Bacillus | no | no | yes | no |
| BV5_Otu000109_Kaistobacter | no | no | yes | yes |
| BV5_Otu000112_Planococcaceae | no | no | yes | no |
| BV5_Otu000113_Dongia | no | no | yes | no |
| BV5_Otu000114_Escherichia | no | no | no | no |
| BV5_Otu000117_Solirubrobacterales | no | no | yes | yes |
| BV5_Otu000118_[Weeksellaceae] | no | no | no | no |
| BV5_Otu000120_Bacillus | no | no | yes | no |
| BV5_Otu000125_Steroidobacter | no | no | yes | no |
| BV5_Otu000141_Ellin6513 | no | no | no | no |
| BV5_Otu000143_Dolo | no | no | yes | no |
| BV5_Otu000144_Rhizobiales | no | no | yes | no |
| BV5_Otu000155_Burkholderia | no | no | no | no |
| BV5_Otu000156_Staphylococcus | no | no | no | no |
| BV5_Otu000194_Micromonospora | no | no | yes | no |
| BV5_Otu000256_Bacillales | no | no | yes | no |
| BV5_Otu000713_Bacteria | no | no | no | no |
| ITS2_Otu00003_Cladosporium | yes | no | yes | yes |
| ITS2_Otu00004_Exophiala | yes | yes | yes | yes |
| ITS2_Otu00005_Cadophora | yes | yes | yes | no |
| ITS2_Otu00006_Fungi | yes | yes | yes | yes |
| ITS2_Otu00008_Boeremia | no | no | no | no |
| ITS2_Otu00009_Didymellaceae | no | no | no | no |
| ITS2_Otu00010_Sclerotiniaceae | yes | no | no | no |
| ITS2_Otu00011_Septoria | yes | no | no | no |
| ITS2_Otu00012_Alternaria | yes | no | no | yes |
| ITS2_Otu00013_Didymellaceae | no | no | no | yes |
| ITS2_Otu00014_Chaetosphaeronema | yes | no | no | yes |
| ITS2_Otu00015_Pseudomassaria | yes | yes | no | no |
| ITS2_Otu00016_Botryotinia | no | no | no | yes |
| ITS2_Otu00017_Helotiales | no | yes | no | yes |
| ITS2_Otu00018_Tetracladium | yes | yes | no | no |
| ITS2_Otu00019_Alternaria | yes | no | no | no |
| ITS2_Otu00020_Podospora | yes | no | no | no |
| ITS2_Otu00021_Dactylonectria | yes | yes | yes | no |
| ITS2_Otu00022_Neocosmospora | yes | yes | no | no |
| ITS2_Otu00023_Cistella | yes | yes | no | no |
| ITS2_Otu00024_Gibberella | no | no | no | no |
| ITS2_Otu00025_Colletotrichum | yes | no | no | no |
| ITS2_Otu00026_Mycena | no | yes | no | no |
| ITS2_Otu00028_Ascomycota | yes | yes | no | no |
| ITS2_Otu00030_Uromyces | yes | no | no | no |
| ITS2_Otu00031_Stemphylium | yes | no | no | no |
| ITS2_Otu00033_Ilyonectria | no | yes | yes | no |
| ITS2_Otu00034_Serendipita | no | yes | no | yes |
| ITS2_Otu00035_Pseudoidium | yes | no | no | no |
| ITS2_Otu00036_Hemimycena | yes | yes | no | no |
| ITS2_Otu00037_Exophiala | no | yes | no | no |
| ITS2_Otu00042_Pleosporales | yes | no | no | no |

| | | | | |
|---|---|---|---|---|
| ITS2_Otu00052_Titaea | yes | no | no | yes |
| ITS2_Otu00054_Ramularia | no | no | no | no |
| ITS2_Otu00058_Colletotrichum | yes | no | no | no |
| ITS2_Otu00063_Pleosporales | no | no | no | no |
| ITS2_Otu00085_Fungi | yes | no | yes | no |
| ITS2_Otu00109_Malassezia | yes | no | no | no |
| ITS2_Otu00232_Fungi | no | no | no | no |
| ITS2_Otu00257_Fungi | no | no | no | no |
| PV9_Otu00002_Pezothrips | yes | no | no | no |
| PV9_Otu00003_Insecta | yes | no | no | no |
| PV9_Otu00004_Frankliniella | yes | no | no | no |
| PV9_Otu00005_Exophiala | yes | yes | yes | yes |
| PV9_Otu00007_Pucciniomycetes | no | no | no | no |
| PV9_Otu00008_Mayetiola | yes | no | no | no |
| PV9_Otu00009_Dothideomycetes | yes | yes | no | yes |
| PV9_Otu00010_Aeolothrips | no | no | no | no |
| PV9_Otu00011_Agaricomycetes | yes | yes | no | no |
| PV9_Otu00012_Agaricomycetes | yes | yes | no | no |
| PV9_Otu00013_Insecta | yes | no | no | no |
| PV9_Otu00014_Metschnikowia | yes | no | no | no |
| PV9_Otu00015_Insecta | no | no | no | no |
| PV9_Otu00016_Agaricomycetes | no | yes | no | no |
| PV9_Otu00018_Aglenchus | yes | yes | no | no |
| PV9_Otu00019_Agaricomycetes | yes | yes | no | no |
| PV9_Otu00021_Insecta | no | no | no | no |
| PV9_Otu00022_Agaricomycetes | yes | yes | no | no |
| PV9_Otu00024_Chromadorea_X | yes | yes | yes | no |
| PV9_Otu00026_Pezizomycotina | yes | yes | no | no |
| PV9_Otu00027_Agaricomycetes | yes | yes | no | no |
| PV9_Otu00028_Dothideomycetes | yes | no | no | no |
| PV9_Otu00030_Arachnida | no | yes | no | no |
| PV9_Otu00032_Arachnida | yes | no | no | no |
| PV9_Otu00033_Arachnida | yes | no | no | no |
| PV9_Otu00034_Pezizomycotina | yes | yes | yes | yes |
| PV9_Otu00035_Paratylenchus | yes | yes | no | no |
| PV9_Otu00038_Insecta | no | no | no | no |
| PV9_Otu00043_Plectus | yes | yes | no | no |
| PV9_Otu00047_Alatospora | no | yes | no | no |
| PV9_Otu00053_Sebacina | no | yes | no | no |
| PV9_Otu00057_Agaricomycetes | no | yes | no | no |
| PV9_Otu00061_Dothideomycetes | no | no | no | yes |
| PV9_Otu00062_Pratylenchus | no | yes | no | no |
| PV9_Otu00064_Aphelenchus | no | yes | no | no |
| PV9_Otu00081_Leotiomycetes | yes | no | no | no |
| PV9_Otu00083_Adineta | no | no | no | no |
| PV9_Otu00084_Phytomyza | no | no | no | no |
| PV9_Otu00101_Exobasidiomycetes | yes | no | no | no |
| PV9_Otu00103_Eukaryota | yes | no | yes | no |
| PV9_Otu00104_Cercomonas | yes | no | yes | yes |
| PV9_Otu00134_Amoebozoa | no | no | yes | yes |
| PV9_Otu00204_Rattus | no | no | no | no |
| PV9_Otu00337_Eukaryota | no | no | yes | no |

**(b)** Shoots

| | Separator | Abundant | Core | Hub |
|---|---|---|---|---|
| BV5_Otu000001_Pantoea | no | yes | yes | no |
| BV5_Otu000003_Pseudomonas | no | yes | yes | no |
| BV5_Otu000004_Phyllobacteriaceae | yes | no | no | no |

| | | | | |
|---|---|---|---|---|
| BV5_Otu000005_Enterobacteriaceae | yes | no | no | no |
| BV5_Otu000006_Ralstonia | no | yes | yes | no |
| BV5_Otu000007_Bacillus | yes | yes | yes | yes |
| BV5_Otu000008_Frankia | no | no | no | no |
| BV5_Otu000009_Phyllobacterium | yes | no | no | yes |
| BV5_Otu000010_Pantoea | no | yes | no | no |
| BV5_Otu000011_Mesorhizobium | no | no | no | yes |
| BV5_Otu000012_Burkholderia | yes | yes | no | no |
| BV5_Otu000013_Burkholderia | yes | yes | no | yes |
| BV5_Otu000014_Agrobacterium | yes | yes | yes | no |
| BV5_Otu000015_Streptomyces | no | no | no | yes |
| BV5_Otu000016_Burkholderia | yes | yes | no | no |
| BV5_Otu000017_Mesorhizobium | no | no | no | no |
| BV5_Otu000018_Comamonadaceae | yes | yes | no | no |
| BV5_Otu000019_Cryptosporangium | yes | no | no | no |
| BV5_Otu000021_Pantoea | no | no | no | no |
| BV5_Otu000022_Methylibium | no | no | no | no |
| BV5_Otu000023_Steroidobacter | no | no | no | no |
| BV5_Otu000024_Xanthomonadaceae | yes | yes | no | yes |
| BV5_Otu000025_Rhodoplanes | no | no | no | yes |
| BV5_Otu000026_Rhizobium | no | no | no | no |
| BV5_Otu000027_Gaiellaceae | no | no | no | yes |
| BV5_Otu000028_Wolbachia | yes | no | no | no |
| BV5_Otu000029_Rhizobiales | no | no | no | yes |
| BV5_Otu000030_0319-7L14 | no | no | no | yes |
| BV5_Otu000032_Pseudomonas | yes | no | no | no |
| BV5_Otu000033_Chryseobacterium | no | no | no | no |
| BV5_Otu000034_Prauserella | yes | no | no | no |
| BV5_Otu000036_Bosea | yes | no | no | yes |
| BV5_Otu000039_Methylibium | no | no | no | no |
| BV5_Otu000040_Sodalis | no | no | no | no |
| BV5_Otu000043_Bradyrhizobiaceae | no | no | no | no |
| BV5_Otu000045_Micromonosporaceae | no | no | no | yes |
| BV5_Otu000047_Methylobacterium | yes | yes | no | no |
| BV5_Otu000049_Kineosporia | no | no | no | no |
| BV5_Otu000052_Methylobacterium | yes | yes | no | yes |
| BV5_Otu000054_Gaiellaceae | no | no | no | yes |
| BV5_Otu000055_Methylobacterium | yes | no | no | yes |
| BV5_Otu000056_Enterobacteriaceae | no | no | no | no |
| BV5_Otu000057_Rhodospirillaceae | no | no | no | yes |
| BV5_Otu000059_Solirubrobacterales | no | no | no | yes |
| BV5_Otu000060_Hyphomicrobium | no | no | no | no |
| BV5_Otu000061_Mycobacterium | no | no | no | yes |
| BV5_Otu000062_Flavobacterium | no | no | no | yes |
| BV5_Otu000063_SC-I-84 | no | no | no | yes |
| BV5_Otu000065_Solirubrobacterales | no | no | no | yes |
| BV5_Otu000068_Gluconobacter | no | no | no | no |
| BV5_Otu000069_Pseudomonas | no | no | no | no |
| BV5_Otu000070_Rubrobacter | yes | no | no | no |
| BV5_Otu000073_Erwinia | yes | no | no | no |
| BV5_Otu000075_Solirubrobacteraceae | no | no | no | yes |
| BV5_Otu000076_Acinetobacter | yes | no | no | no |
| BV5_Otu000077_Solirubrobacterales | no | no | no | yes |
| BV5_Otu000078_Sinobacteraceae | no | no | no | yes |
| BV5_Otu000079_Buchnera | no | no | no | no |
| BV5_Otu000080_Pedomicrobium | no | no | no | yes |
| BV5_Otu000081_SC-I-84 | no | no | no | yes |
| BV5_Otu000084_Dongia | no | no | no | no |

| | | | | |
|---|---|---|---|---|
| BV5_Otu000086_Burkholderiaceae | yes | no | no | no |
| BV5_Otu000089_Propionibacterium | yes | no | yes | no |
| BV5_Otu000091_Patulibacteraceae | no | no | no | no |
| BV5_Otu000092_0319-7L14 | no | no | no | yes |
| BV5_Otu000094_Caulobacter | no | no | no | no |
| BV5_Otu000095_Sphingomonas | yes | no | yes | no |
| BV5_Otu000096_Bradyrhizobium | no | no | no | no |
| BV5_Otu000097_Gaiellaceae | no | no | no | yes |
| BV5_Otu000098_Mycobacterium | no | no | no | no |
| BV5_Otu000102_Solibacillus | no | no | no | yes |
| BV5_Otu000105_Labrys | no | no | no | no |
| BV5_Otu000106_Bacillus | no | no | no | yes |
| BV5_Otu000109_Kaistobacter | no | no | no | yes |
| BV5_Otu000112_Planococcaceae | no | no | no | yes |
| BV5_Otu000113_Dongia | no | no | no | no |
| BV5_Otu000114_Escherichia | yes | no | no | no |
| BV5_Otu000117_Solirubrobacterales | no | no | no | yes |
| BV5_Otu000118_[Weeksellaceae] | yes | no | no | no |
| BV5_Otu000120_Bacillus | no | no | no | yes |
| BV5_Otu000125_Steroidobacter | no | no | no | no |
| BV5_Otu000141_Ellin6513 | no | no | no | no |
| BV5_Otu000143_Dolo | no | no | no | no |
| BV5_Otu000144_Rhizobiales | no | no | no | no |
| BV5_Otu000155_Burkholderia | no | no | no | no |
| BV5_Otu000156_Staphylococcus | yes | no | no | no |
| BV5_Otu000194_Micromonospora | no | no | no | yes |
| BV5_Otu000256_Bacillales | no | no | no | yes |
| BV5_Otu000713_Bacteria | no | no | no | no |
| ITS2_Otu00003_Cladosporium | no | yes | yes | yes |
| ITS2_Otu00004_Exophiala | yes | no | no | no |
| ITS2_Otu00005_Cadophora | yes | no | no | no |
| ITS2_Otu00006_Fungi | no | yes | yes | no |
| ITS2_Otu00008_Boeremia | yes | yes | no | yes |
| ITS2_Otu00009_Didymellaceae | yes | yes | no | yes |
| ITS2_Otu00010_Sclerotiniaceae | yes | no | no | no |
| ITS2_Otu00011_Septoria | yes | yes | no | yes |
| ITS2_Otu00012_Alternaria | no | yes | no | yes |
| ITS2_Otu00013_Didymellaceae | yes | yes | no | no |
| ITS2_Otu00014_Chaetosphaeronema | yes | yes | no | yes |
| ITS2_Otu00015_Pseudomassaria | yes | no | no | no |
| ITS2_Otu00016_Botryotinia | yes | no | no | no |
| ITS2_Otu00017_Helotiales | yes | no | no | no |
| ITS2_Otu00018_Tetracladium | yes | no | no | no |
| ITS2_Otu00019_Alternaria | yes | yes | yes | yes |
| ITS2_Otu00020_Podospora | yes | yes | no | no |
| ITS2_Otu00021_Dactylonectria | no | no | no | no |
| ITS2_Otu00022_Neocosmospora | no | no | no | no |
| ITS2_Otu00023_Cistella | no | no | no | no |
| ITS2_Otu00024_Gibberella | no | yes | no | yes |
| ITS2_Otu00025_Colletotrichum | yes | yes | no | yes |
| ITS2_Otu00026_Mycena | no | no | no | no |
| ITS2_Otu00028_Ascomycota | no | no | no | no |
| ITS2_Otu00030_Uromyces | yes | yes | no | no |
| ITS2_Otu00031_Stemphylium | yes | yes | no | no |
| ITS2_Otu00033_Ilyonectria | no | no | no | no |
| ITS2_Otu00034_Serendipita | no | no | no | no |
| ITS2_Otu00035_Pseudoidium | yes | yes | no | no |
| ITS2_Otu00036_Hemimycena | no | no | no | no |

| | | | | |
|---|---|---|---|---|
| ITS2_Otu00037_Exophiala | no | no | no | no |
| ITS2_Otu00042_Pleosporales | no | no | no | no |
| ITS2_Otu00052_Titaea | no | no | no | no |
| ITS2_Otu00054_Ramularia | yes | yes | no | no |
| ITS2_Otu00058_Colletotrichum | yes | yes | no | yes |
| ITS2_Otu00063_Pleosporales | yes | yes | no | yes |
| ITS2_Otu00085_Fungi | yes | no | yes | no |
| ITS2_Otu00109_Malassezia | no | no | no | no |
| ITS2_Otu00232_Fungi | no | no | no | no |
| ITS2_Otu00257_Fungi | no | no | no | no |
| PV9_Otu00002_Pezothrips | no | yes | yes | no |
| PV9_Otu00003_Insecta | yes | no | no | no |
| PV9_Otu00004_Frankliniella | no | yes | no | no |
| PV9_Otu00005_Exophiala | yes | no | no | no |
| PV9_Otu00007_Pucciniomycetes | yes | yes | no | no |
| PV9_Otu00008_Mayetiola | yes | yes | no | no |
| PV9_Otu00009_Dothideomycetes | yes | yes | yes | yes |
| PV9_Otu00010_Aeolothrips | no | no | no | no |
| PV9_Otu00011_Agaricomycetes | yes | no | no | no |
| PV9_Otu00012_Agaricomycetes | no | no | no | yes |
| PV9_Otu00013_Insecta | no | yes | no | no |
| PV9_Otu00014_Metschnikowia | yes | no | no | no |
| PV9_Otu00015_Insecta | no | yes | no | no |
| PV9_Otu00016_Agaricomycetes | no | no | no | no |
| PV9_Otu00018_Aglenchus | no | no | no | no |
| PV9_Otu00019_Agaricomycetes | yes | no | no | no |
| PV9_Otu00021_Insecta | no | no | no | no |
| PV9_Otu00022_Agaricomycetes | no | no | no | no |
| PV9_Otu00024_Chromadorea_X | yes | no | no | no |
| PV9_Otu00026_Pezizomycotina | no | no | no | no |
| PV9_Otu00027_Agaricomycetes | no | no | no | no |
| PV9_Otu00028_Dothideomycetes | no | no | no | yes |
| PV9_Otu00030_Arachnida | no | no | no | no |
| PV9_Otu00032_Arachnida | yes | yes | no | no |
| PV9_Otu00033_Arachnida | no | no | no | no |
| PV9_Otu00034_Pezizomycotina | yes | no | no | no |
| PV9_Otu00035_Paratylenchus | no | no | no | no |
| PV9_Otu00038_Insecta | yes | yes | no | no |
| PV9_Otu00043_Plectus | no | no | no | no |
| PV9_Otu00047_Alatospora | no | no | no | no |
| PV9_Otu00053_Sebacina | no | no | no | no |
| PV9_Otu00057_Agaricomycetes | no | no | no | no |
| PV9_Otu00061_Dothideomycetes | yes | no | no | yes |
| PV9_Otu00062_Pratylenchus | no | no | no | no |
| PV9_Otu00064_Aphelenchus | no | no | no | no |
| PV9_Otu00081_Leotiomycetes | yes | yes | no | no |
| PV9_Otu00083_Adineta | yes | no | no | yes |
| PV9_Otu00084_Phytomyza | yes | yes | no | no |
| PV9_Otu00101_Exobasidiomycetes | yes | no | no | no |
| PV9_Otu00103_Eukaryota | yes | no | yes | no |
| PV9_Otu00104_Cercomonas | no | no | no | no |
| PV9_Otu00134_Amoebozoa | no | no | no | no |
| PV9_Otu00204_Rattus | no | no | no | no |
| PV9_Otu00337_Eukaryota | no | no | yes | no |

**(c)** Flowers

| | Separator | Abundant | Core | Hub |
|---|---|---|---|---|
| BV5_Otu000001_Pantoea | no | yes | no | no |

| | | | | |
|---|---|---|---|---|
| BV5_Otu000003_Pseudomonas | no | yes | yes | yes |
| BV5_Otu000004_Phyllobacteriaceae | no | no | no | yes |
| BV5_Otu000005_Enterobacteriaceae | yes | yes | no | no |
| BV5_Otu000006_Ralstonia | yes | yes | no | yes |
| BV5_Otu000007_Bacillus | yes | no | no | yes |
| BV5_Otu000008_Frankia | no | no | no | no |
| BV5_Otu000009_Phyllobacterium | no | no | no | yes |
| BV5_Otu000010_Pantoea | no | yes | no | yes |
| BV5_Otu000011_Mesorhizobium | no | no | no | yes |
| BV5_Otu000012_Burkholderia | yes | no | no | yes |
| BV5_Otu000013_Burkholderia | yes | no | no | yes |
| BV5_Otu000014_Agrobacterium | yes | no | no | no |
| BV5_Otu000015_Streptomyces | no | no | no | yes |
| BV5_Otu000016_Burkholderia | yes | no | no | yes |
| BV5_Otu000017_Mesorhizobium | no | no | no | yes |
| BV5_Otu000018_Comamonadaceae | no | no | no | no |
| BV5_Otu000019_Cryptosporangium | no | no | no | no |
| BV5_Otu000021_Pantoea | no | yes | no | yes |
| BV5_Otu000022_Methylibium | no | no | no | no |
| BV5_Otu000023_Steroidobacter | no | no | no | no |
| BV5_Otu000024_Xanthomonadaceae | yes | no | no | no |
| BV5_Otu000025_Rhodoplanes | no | no | no | yes |
| BV5_Otu000026_Rhizobium | no | no | no | yes |
| BV5_Otu000027_Gaiellaceae | no | no | no | yes |
| BV5_Otu000028_Wolbachia | yes | yes | no | no |
| BV5_Otu000029_Rhizobiales | no | no | no | yes |
| BV5_Otu000030_0319-7L14 | no | no | no | yes |
| BV5_Otu000032_Pseudomonas | yes | yes | no | yes |
| BV5_Otu000033_Chryseobacterium | no | no | no | no |
| BV5_Otu000034_Prauserella | yes | no | no | yes |
| BV5_Otu000036_Bosea | yes | no | no | no |
| BV5_Otu000039_Methylibium | no | no | no | no |
| BV5_Otu000040_Sodalis | no | no | no | no |
| BV5_Otu000043_Bradyrhizobiaceae | no | no | no | yes |
| BV5_Otu000045_Micromonosporaceae | no | no | no | no |
| BV5_Otu000047_Methylobacterium | yes | no | no | no |
| BV5_Otu000049_Kineosporia | no | no | no | no |
| BV5_Otu000052_Methylobacterium | no | no | no | yes |
| BV5_Otu000054_Gaiellaceae | no | no | no | yes |
| BV5_Otu000055_Methylobacterium | no | no | no | yes |
| BV5_Otu000056_Enterobacteriaceae | yes | yes | no | no |
| BV5_Otu000057_Rhodospirillaceae | no | no | no | yes |
| BV5_Otu000059_Solirubrobacterales | no | no | no | yes |
| BV5_Otu000060_Hyphomicrobium | no | no | no | no |
| BV5_Otu000061_Mycobacterium | no | no | no | yes |
| BV5_Otu000062_Flavobacterium | no | no | no | no |
| BV5_Otu000063_SC-I-84 | no | no | no | yes |
| BV5_Otu000065_Solirubrobacterales | no | no | no | yes |
| BV5_Otu000068_Gluconobacter | no | yes | no | no |
| BV5_Otu000069_Pseudomonas | no | yes | no | no |
| BV5_Otu000070_Rubrobacter | no | no | no | yes |
| BV5_Otu000073_Erwinia | no | no | no | yes |
| BV5_Otu000075_Solirubrobacteraceae | no | no | no | yes |
| BV5_Otu000076_Acinetobacter | yes | yes | no | no |
| BV5_Otu000077_Solirubrobacterales | no | no | no | yes |
| BV5_Otu000078_Sinobacteraceae | no | no | no | yes |
| BV5_Otu000079_Buchnera | no | yes | no | no |
| BV5_Otu000080_Pedomicrobium | no | no | no | no |

| | | | | |
|---|---|---|---|---|
| BV5_Otu000081_SC-I-84 | no | no | no | yes |
| BV5_Otu000084_Dongia | no | no | no | no |
| BV5_Otu000086_Burkholderiaceae | no | no | no | yes |
| BV5_Otu000089_Propionibacterium | yes | no | no | yes |
| BV5_Otu000091_Patulibacteraceae | no | no | no | no |
| BV5_Otu000092_0319-7L14 | no | no | no | yes |
| BV5_Otu000094_Caulobacter | no | no | no | no |
| BV5_Otu000095_Sphingomonas | no | no | no | yes |
| BV5_Otu000096_Bradyrhizobium | no | no | no | yes |
| BV5_Otu000097_Gaiellaceae | no | no | no | yes |
| BV5_Otu000098_Mycobacterium | no | no | no | yes |
| BV5_Otu000102_Solibacillus | no | no | no | no |
| BV5_Otu000105_Labrys | no | no | no | no |
| BV5_Otu000106_Bacillus | no | no | no | no |
| BV5_Otu000109_Kaistobacter | no | no | no | yes |
| BV5_Otu000112_Planococcaceae | no | no | no | no |
| BV5_Otu000113_Dongia | no | no | no | no |
| BV5_Otu000114_Escherichia | yes | no | no | yes |
| BV5_Otu000117_Solirubrobacterales | no | no | no | no |
| BV5_Otu000118_[Weeksellaceae] | no | no | no | yes |
| BV5_Otu000120_Bacillus | no | no | no | no |
| BV5_Otu000125_Steroidobacter | no | no | no | no |
| BV5_Otu000141_Ellin6513 | no | no | no | yes |
| BV5_Otu000143_Dolo | no | no | no | no |
| BV5_Otu000144_Rhizobiales | no | no | no | no |
| BV5_Otu000155_Burkholderia | no | no | no | no |
| BV5_Otu000156_Staphylococcus | yes | no | no | no |
| BV5_Otu000194_Micromonospora | no | no | no | yes |
| BV5_Otu000256_Bacillales | no | no | no | no |
| BV5_Otu000713_Bacteria | no | no | no | no |
| ITS2_Otu00003_Cladosporium | yes | yes | yes | yes |
| ITS2_Otu00004_Exophiala | no | no | no | no |
| ITS2_Otu00005_Cadophora | no | no | no | no |
| ITS2_Otu00006_Fungi | no | yes | yes | yes |
| ITS2_Otu00008_Boeremia | yes | no | no | yes |
| ITS2_Otu00009_Didymellaceae | yes | yes | no | yes |
| ITS2_Otu00010_Sclerotiniaceae | yes | yes | no | yes |
| ITS2_Otu00011_Septoria | yes | no | no | no |
| ITS2_Otu00012_Alternaria | yes | yes | no | yes |
| ITS2_Otu00013_Didymellaceae | yes | yes | no | yes |
| ITS2_Otu00014_Chaetosphaeronema | yes | no | no | no |
| ITS2_Otu00015_Pseudomassaria | no | no | no | no |
| ITS2_Otu00016_Botryotinia | yes | yes | no | yes |
| ITS2_Otu00017_Helotiales | no | no | no | no |
| ITS2_Otu00018_Tetracladium | no | no | no | no |
| ITS2_Otu00019_Alternaria | no | yes | no | yes |
| ITS2_Otu00020_Podospora | yes | no | no | no |
| ITS2_Otu00021_Dactylonectria | no | no | no | no |
| ITS2_Otu00022_Neocosmospora | no | no | no | no |
| ITS2_Otu00023_Cistella | no | no | no | no |
| ITS2_Otu00024_Gibberella | no | no | no | no |
| ITS2_Otu00025_Colletotrichum | yes | no | no | no |
| ITS2_Otu00026_Mycena | no | no | no | no |
| ITS2_Otu00028_Ascomycota | no | no | no | no |
| ITS2_Otu00030_Uromyces | no | no | no | no |
| ITS2_Otu00031_Stemphylium | yes | no | no | no |
| ITS2_Otu00033_Ilyonectria | no | no | no | no |
| ITS2_Otu00034_Serendipita | no | no | no | no |

| | | | | |
|---|---|---|---|---|
| ITS2_Otu00035_Pseudoidium | no | no | no | no |
| ITS2_Otu00036_Hemimycena | no | no | no | no |
| ITS2_Otu00037_Exophiala | no | no | no | no |
| ITS2_Otu00042_Pleosporales | no | no | no | no |
| ITS2_Otu00052_Titaea | no | no | no | no |
| ITS2_Otu00054_Ramularia | yes | no | no | no |
| ITS2_Otu00058_Colletotrichum | no | no | no | no |
| ITS2_Otu00063_Pleosporales | no | no | no | no |
| ITS2_Otu00085_Fungi | no | no | yes | no |
| ITS2_Otu00109_Malassezia | no | no | no | no |
| ITS2_Otu00232_Fungi | no | no | no | no |
| ITS2_Otu00257_Fungi | no | no | no | no |
| PV9_Otu00002_Pezothrips | yes | yes | yes | no |
| PV9_Otu00003_Insecta | yes | no | no | no |
| PV9_Otu00004_Frankliniella | yes | yes | no | no |
| PV9_Otu00005_Exophiala | no | no | no | no |
| PV9_Otu00007_Pucciniomycetes | no | no | no | no |
| PV9_Otu00008_Mayetiola | yes | yes | no | no |
| PV9_Otu00009_Dothideomycetes | yes | yes | no | yes |
| PV9_Otu00010_Aeolothrips | yes | yes | no | no |
| PV9_Otu00011_Agaricomycetes | no | no | no | no |
| PV9_Otu00012_Agaricomycetes | no | no | no | no |
| PV9_Otu00013_Insecta | no | yes | no | no |
| PV9_Otu00014_Metschnikowia | yes | yes | no | yes |
| PV9_Otu00015_Insecta | yes | yes | no | no |
| PV9_Otu00016_Agaricomycetes | no | no | no | no |
| PV9_Otu00018_Aglenchus | no | no | no | no |
| PV9_Otu00019_Agaricomycetes | no | no | no | no |
| PV9_Otu00021_Insecta | yes | yes | no | no |
| PV9_Otu00022_Agaricomycetes | no | no | no | no |
| PV9_Otu00024_Chromadorea_X | no | no | no | no |
| PV9_Otu00026_Pezizomycotina | no | no | no | no |
| PV9_Otu00027_Agaricomycetes | no | no | no | no |
| PV9_Otu00028_Dothideomycetes | no | yes | no | yes |
| PV9_Otu00030_Arachnida | no | no | no | no |
| PV9_Otu00032_Arachnida | no | no | no | no |
| PV9_Otu00033_Arachnida | yes | yes | no | no |
| PV9_Otu00034_Pezizomycotina | no | no | no | no |
| PV9_Otu00035_Paratylenchus | no | no | no | no |
| PV9_Otu00038_Insecta | yes | no | no | no |
| PV9_Otu00043_Plectus | no | no | no | no |
| PV9_Otu00047_Alatospora | no | no | no | no |
| PV9_Otu00053_Sebacina | no | no | no | no |
| PV9_Otu00057_Agaricomycetes | no | no | no | no |
| PV9_Otu00061_Dothideomycetes | yes | no | no | no |
| PV9_Otu00062_Pratylenchus | no | no | no | no |
| PV9_Otu00064_Aphelenchus | no | no | no | no |
| PV9_Otu00081_Leotiomycetes | no | no | no | no |
| PV9_Otu00083_Adineta | no | no | no | no |
| PV9_Otu00084_Phytomyza | no | no | no | no |
| PV9_Otu00101_Exobasidiomycetes | no | no | no | no |
| PV9_Otu00103_Eukaryota | yes | no | yes | no |
| PV9_Otu00104_Cercomonas | no | no | no | no |
| PV9_Otu00134_Amoebozoa | no | no | no | no |
| PV9_Otu00204_Rattus | no | no | no | no |
| PV9_Otu00337_Eukaryota | no | no | yes | no |

**(d)** Seeds

| | Separator | Abundant | Core | Hub |
|---|---|---|---|---|
| BV5_Otu000001_Pantoea | yes | yes | no | no |
| BV5_Otu000003_Pseudomonas | yes | yes | yes | yes |
| BV5_Otu000004_Phyllobacteriaceae | yes | no | no | yes |
| BV5_Otu000005_Enterobacteriaceae | no | yes | no | no |
| BV5_Otu000006_Ralstonia | yes | yes | yes | yes |
| BV5_Otu000007_Bacillus | yes | no | no | yes |
| BV5_Otu000008_Frankia | yes | no | no | no |
| BV5_Otu000009_Phyllobacterium | yes | no | no | no |
| BV5_Otu000010_Pantoea | no | yes | no | no |
| BV5_Otu000011_Mesorhizobium | yes | no | no | no |
| BV5_Otu000012_Burkholderia | yes | yes | no | yes |
| BV5_Otu000013_Burkholderia | yes | yes | no | yes |
| BV5_Otu000014_Agrobacterium | no | no | no | yes |
| BV5_Otu000015_Streptomyces | no | no | no | no |
| BV5_Otu000016_Burkholderia | yes | yes | no | yes |
| BV5_Otu000017_Mesorhizobium | yes | no | no | yes |
| BV5_Otu000018_Comamonadaceae | yes | yes | no | no |
| BV5_Otu000019_Cryptosporangium | yes | no | no | no |
| BV5_Otu000021_Pantoea | yes | yes | no | no |
| BV5_Otu000022_Methylibium | no | no | no | no |
| BV5_Otu000023_Steroidobacter | no | no | no | no |
| BV5_Otu000024_Xanthomonadaceae | yes | no | no | no |
| BV5_Otu000025_Rhodoplanes | no | no | no | no |
| BV5_Otu000026_Rhizobium | yes | no | no | no |
| BV5_Otu000027_Gaiellaceae | no | no | no | no |
| BV5_Otu000028_Wolbachia | yes | yes | no | no |
| BV5_Otu000029_Rhizobiales | no | no | no | no |
| BV5_Otu000030_0319-7L14 | no | no | no | no |
| BV5_Otu000032_Pseudomonas | yes | no | no | yes |
| BV5_Otu000033_Chryseobacterium | no | no | no | no |
| BV5_Otu000034_Prauserella | yes | yes | no | yes |
| BV5_Otu000036_Bosea | yes | no | no | no |
| BV5_Otu000039_Methylibium | no | no | no | no |
| BV5_Otu000040_Sodalis | yes | yes | no | no |
| BV5_Otu000043_Bradyrhizobiaceae | no | no | no | yes |
| BV5_Otu000045_Micromonosporaceae | no | no | no | no |
| BV5_Otu000047_Methylobacterium | no | no | no | no |
| BV5_Otu000049_Kineosporia | yes | no | no | no |
| BV5_Otu000052_Methylobacterium | yes | no | no | yes |
| BV5_Otu000054_Gaiellaceae | no | no | no | no |
| BV5_Otu000055_Methylobacterium | no | no | no | no |
| BV5_Otu000056_Enterobacteriaceae | no | no | no | no |
| BV5_Otu000057_Rhodospirillaceae | no | no | no | yes |
| BV5_Otu000059_Solirubrobacterales | no | no | no | no |
| BV5_Otu000060_Hyphomicrobium | no | no | no | no |
| BV5_Otu000061_Mycobacterium | no | no | no | no |
| BV5_Otu000062_Flavobacterium | no | no | no | no |
| BV5_Otu000063_SC-I-84 | no | no | no | no |
| BV5_Otu000065_Solirubrobacterales | no | no | no | no |
| BV5_Otu000068_Gluconobacter | no | no | no | no |
| BV5_Otu000069_Pseudomonas | no | no | no | no |
| BV5_Otu000070_Rubrobacter | yes | no | no | yes |
| BV5_Otu000073_Erwinia | yes | no | no | yes |
| BV5_Otu000075_Solirubrobacteraceae | no | no | no | no |
| BV5_Otu000076_Acinetobacter | no | no | no | no |
| BV5_Otu000077_Solirubrobacterales | no | no | no | no |
| BV5_Otu000078_Sinobacteraceae | no | no | no | no |

| | | | | |
|---|---|---|---|---|
| BV5_Otu000079_Buchnera | no | no | no | no |
| BV5_Otu000080_Pedomicrobium | no | no | no | no |
| BV5_Otu000081_SC-I-84 | no | no | no | no |
| BV5_Otu000084_Dongia | no | no | no | no |
| BV5_Otu000086_Burkholderiaceae | yes | no | no | yes |
| BV5_Otu000089_Propionibacterium | yes | no | yes | yes |
| BV5_Otu000091_Patulibacteraceae | no | no | no | no |
| BV5_Otu000092_0319-7L14 | no | no | no | no |
| BV5_Otu000094_Caulobacter | no | no | no | no |
| BV5_Otu000095_Sphingomonas | yes | no | no | no |
| BV5_Otu000096_Bradyrhizobium | no | no | no | yes |
| BV5_Otu000097_Gaiellaceae | no | no | no | no |
| BV5_Otu000098_Mycobacterium | no | no | no | no |
| BV5_Otu000102_Solibacillus | no | no | no | no |
| BV5_Otu000105_Labrys | no | no | no | no |
| BV5_Otu000106_Bacillus | no | no | no | no |
| BV5_Otu000109_Kaistobacter | no | no | no | no |
| BV5_Otu000112_Planococcaceae | no | no | no | no |
| BV5_Otu000113_Dongia | no | no | no | no |
| BV5_Otu000114_Escherichia | no | no | no | yes |
| BV5_Otu000117_Solirubrobacterales | no | no | no | no |
| BV5_Otu000118_[Weeksellaceae] | yes | no | no | yes |
| BV5_Otu000120_Bacillus | no | no | no | no |
| BV5_Otu000125_Steroidobacter | no | no | no | no |
| BV5_Otu000141_Ellin6513 | yes | no | no | yes |
| BV5_Otu000143_Dolo | no | no | no | no |
| BV5_Otu000144_Rhizobiales | no | no | no | no |
| BV5_Otu000155_Burkholderia | yes | no | no | yes |
| BV5_Otu000156_Staphylococcus | yes | no | no | yes |
| BV5_Otu000194_Micromonospora | no | no | no | no |
| BV5_Otu000256_Bacillales | no | no | no | no |
| BV5_Otu000713_Bacteria | yes | no | no | yes |
| ITS2_Otu00003_Cladosporium | yes | yes | yes | no |
| ITS2_Otu00004_Exophiala | yes | no | no | no |
| ITS2_Otu00005_Cadophora | yes | no | no | no |
| ITS2_Otu00006_Fungi | yes | yes | yes | yes |
| ITS2_Otu00008_Boeremia | yes | yes | no | no |
| ITS2_Otu00009_Didymellaceae | yes | no | no | no |
| ITS2_Otu00010_Sclerotiniaceae | yes | yes | no | no |
| ITS2_Otu00011_Septoria | yes | no | no | no |
| ITS2_Otu00012_Alternaria | no | yes | no | no |
| ITS2_Otu00013_Didymellaceae | yes | yes | no | no |
| ITS2_Otu00014_Chaetosphaeronema | yes | no | no | no |
| ITS2_Otu00015_Pseudomassaria | yes | no | no | no |
| ITS2_Otu00016_Botryotinia | no | no | no | no |
| ITS2_Otu00017_Helotiales | no | no | no | no |
| ITS2_Otu00018_Tetracladium | no | no | no | no |
| ITS2_Otu00019_Alternaria | no | yes | no | no |
| ITS2_Otu00020_Podospora | yes | no | no | no |
| ITS2_Otu00021_Dactylonectria | no | no | no | no |
| ITS2_Otu00022_Neocosmospora | no | no | no | no |
| ITS2_Otu00023_Cistella | no | no | no | no |
| ITS2_Otu00024_Gibberella | yes | yes | no | no |
| ITS2_Otu00025_Colletotrichum | yes | no | no | no |
| ITS2_Otu00026_Mycena | no | no | no | no |
| ITS2_Otu00028_Ascomycota | yes | no | no | no |
| ITS2_Otu00030_Uromyces | no | no | no | no |
| ITS2_Otu00031_Stemphylium | yes | no | no | no |

| | | | | |
|---|---|---|---|---|
| ITS2_Otu00033_Ilyonectria | no | no | no | no |
| ITS2_Otu00034_Serendipita | no | no | no | no |
| ITS2_Otu00035_Pseudoidium | no | no | no | no |
| ITS2_Otu00036_Hemimycena | no | no | no | no |
| ITS2_Otu00037_Exophiala | no | no | no | no |
| ITS2_Otu00042_Pleosporales | no | no | no | no |
| ITS2_Otu00052_Titaea | no | no | no | no |
| ITS2_Otu00054_Ramularia | no | no | no | no |
| ITS2_Otu00058_Colletotrichum | no | no | no | no |
| ITS2_Otu00063_Pleosporales | no | no | no | no |
| ITS2_Otu00085_Fungi | yes | yes | yes | no |
| ITS2_Otu00109_Malassezia | yes | yes | no | no |
| ITS2_Otu00232_Fungi | yes | no | no | yes |
| ITS2_Otu00257_Fungi | yes | no | no | yes |
| PV9_Otu00002_Pezothrips | yes | no | no | no |
| PV9_Otu00003_Insecta | yes | yes | no | no |
| PV9_Otu00004_Frankliniella | yes | no | no | no |
| PV9_Otu00005_Exophiala | yes | no | no | no |
| PV9_Otu00007_Pucciniomycetes | yes | no | no | no |
| PV9_Otu00008_Mayetiola | yes | no | no | no |
| PV9_Otu00009_Dothideomycetes | yes | no | no | no |
| PV9_Otu00010_Aeolothrips | yes | no | no | no |
| PV9_Otu00011_Agaricomycetes | no | no | no | no |
| PV9_Otu00012_Agaricomycetes | no | no | no | no |
| PV9_Otu00013_Insecta | yes | no | no | no |
| PV9_Otu00014_Metschnikowia | yes | no | no | no |
| PV9_Otu00015_Insecta | yes | yes | no | no |
| PV9_Otu00016_Agaricomycetes | yes | no | no | no |
| PV9_Otu00018_Aglenchus | yes | no | no | no |
| PV9_Otu00019_Agaricomycetes | yes | no | no | no |
| PV9_Otu00021_Insecta | yes | no | no | no |
| PV9_Otu00022_Agaricomycetes | yes | no | no | no |
| PV9_Otu00024_Chromadorea_X | yes | no | no | no |
| PV9_Otu00026_Pezizomycotina | yes | no | no | no |
| PV9_Otu00027_Agaricomycetes | yes | no | no | no |
| PV9_Otu00028_Dothideomycetes | yes | no | no | no |
| PV9_Otu00030_Arachnida | no | no | no | no |
| PV9_Otu00032_Arachnida | yes | no | no | no |
| PV9_Otu00033_Arachnida | no | no | no | no |
| PV9_Otu00034_Pezizomycotina | no | no | no | no |
| PV9_Otu00035_Paratylenchus | yes | no | no | no |
| PV9_Otu00038_Insecta | no | no | no | no |
| PV9_Otu00043_Plectus | no | no | no | no |
| PV9_Otu00047_Alatospora | no | no | no | no |
| PV9_Otu00053_Sebacina | no | no | no | no |
| PV9_Otu00057_Agaricomycetes | no | no | no | no |
| PV9_Otu00061_Dothideomycetes | no | no | no | no |
| PV9_Otu00062_Pratylenchus | no | no | no | no |
| PV9_Otu00064_Aphelenchus | no | no | no | yes |
| PV9_Otu00081_Leotiomycetes | yes | no | no | no |
| PV9_Otu00083_Adineta | yes | no | no | no |
| PV9_Otu00084_Phytomyza | yes | no | no | no |
| PV9_Otu00101_Exobasidiomycetes | yes | yes | no | yes |
| PV9_Otu00103_Eukaryota | yes | no | yes | no |
| PV9_Otu00104_Cercomonas | yes | no | no | no |
| PV9_Otu00134_Amoebozoa | no | no | no | no |
| PV9_Otu00204_Rattus | yes | no | no | yes |
| PV9_Otu00337_Eukaryota | yes | no | yes | no |

SUPPLEMENTARY METHODS

**Supplementary Method 1. Sequence data processing using Mothur.**

We processed amplicon sequence data of microbial 16S rRNA, ITS2, and 18S rRNA using Mothur as described in Almario *et al*. (1, 2). In brief, processing of amplicon reads include forming contigs by pairing single-end reads (*make.contigs*), quality filtering for paired reads that are 100-600 bases long with at least 5 bases overlap (*screen.seqs*), demultiplexing (*rename.seqs*), dereplication (*unique.seqs, count.seqs*), detecting and removing chimera using VSEARCH (*chimera.vsearch, remove.seqs*), classifying sequences (*classify.seqs*), OTU clustering at 97% sequence similarity threshold using dgc method (*cluster*), abundance filtering for OTUs with more than 50 reads (*split.abund*), classifying OTUs (*classify.otu*), creating OTU tables (*make.shared*), removing OTUs classified as chloroplast, mitochondria, *Arabidopsis*, Embryophyceae, unknown, and PhiX sequences (*remove.lineage*), and identifying OTU representative sequences based on abundance (*get.oturep*).

**Supplementary Method 2. Identification of hub microbes in *L. corniculatus*.**

To determine hub bacteria, fungi, and eukaryotes in plant organs, correlation networks for each plant organ were computed using SparCC algorithm, as described in Almario *et al*. (1, 3). In brief, OTU absolute count tables with low abundance OTUs removed were used as input for FastSpar, a parallelizable implementation of SparCC algorithm, to calculate correlations (4). P-values were calculated from 1,000 bootstraps and resulting correlations with $P$<0.001 were used for constructing networks.

**Supplementary Method 3. Transmission of microbes in *L. corniculatus***.

To determine how *L. corniculatus* organ microbiomes can be linked and influenced by dispersal, we used Sankey diagrams to visualize potential flow of microbes across the soil and plant organs. Nodes of the diagrams represent different potential microbial sources (*i.e.* soil, plant organs, and environment/others) and arcs correspond to the number of OTUs shared between nodes.

To determine potential origins of organ-associated microbial communities, we used FEAST (Fast Expectation-mAximization microbial Source Tracking) to estimate contribution of potential microbial sources, such as soil, the different plant compartments, or the environment, to each plant organ microbiome (5). In the FEAST analysis we used multidirectional approach where microbiomes from plant organs or soil samples can be both sources or sinks. We implemented FEAST in R using default parameters for 1,000 iterations and we tested sinks with the same group of sources (*i.e.* all source microbiomes throughout seven sites for four years). In FEAST a proportion of the sink microbiomes is also potentially transmitted from "unknown sources", which can be other microbial sources that are not the assigned source in the analysis.

To statistically predict potential priority effects phenomena from roots, shoots, flowers, to seeds in *L. corniculatus*, we identified taxa of interest that are potentially involved in such phenomena in plant organs, as described in Debray *et al*. (6). We examined abundant, core, hub, or machine learning-predicted OTUs if changes in their relative abundance in plant organs correlate with changes in microbial community structure. We used PCoA of Bray-Curtis dissimilarities of plant organ microbial communities and PERMANOVA to assess the influence of these OTUs on community structure when they are abundant or rare (*i.e.* abundant OTUs: Relative abundance in each plant organ (RA) >= 0.01; intermediate OTUs: RA >= 0.001 and < 0.01; rare OTUs: RA < 0.001).
